# Modulation of A2aR Oligomerisation by Conformational State and PIP_2_ Interactions Revealed by MD Simulations and Markov Models

**DOI:** 10.1101/2020.06.24.168260

**Authors:** Wanling Song, Anna L. Duncan, Mark S.P. Sansom

## Abstract

G protein-coupled receptors (GPCRs) play key roles in cellular signalling. GPCRs are suggested to form dimers and higher order oligomers in response to activation. However, we do not fully understand GPCR activation at larger scales and in an *in vivo* context. We have characterised oligomeric configurations of the adenosine 2a receptor (A2aR) by combining large-scale molecular dynamics simulations with Markov state models. Receptor activation results in enhanced oligomerisation, more diverse oligomer populations, and a more connected oligomerisation network. The active state conformation of the A2aR shifts protein-protein association interfaces to those involving intracellular loop ICL3 and transmembrane helix TM6. Binding of PIP_2_ to A2aR stabilises protein-protein interactions via PIP_2_-mediated association interfaces. These results indicate that A2aR oligomerisation is responsive to the local membrane lipid environment. This in turn suggests a modulatory effect on A2aR whereby a given oligomerisation profile favours the dynamic formation of specific supra-molecular signalling complexes.

## Introduction

G Protein-Coupled Receptors (GPCRs) are the largest superfamily in the human genome. They are major drug targets due to involvement in many physiological processes. The canonical model of GPCR activation involves the formation of a ternary structure of a receptor, an agonist and a signalling partner (*1*). However, this paradigm may require a revision given increasing evidence of the existence of GPCR oligomerisation and its biological importance (*2*). For example, single-molecule imaging has revealed that a range of GPCRs (e.g. the dopamine D_2_ receptor and the M_2_ muscarinic receptor) can form metastable oligomers in living cells with lifetimes ranging from sub-milliseconds to seconds (*3, 4*). The degree of oligomerisation varies among receptors. For example, at moderate expression levels, β_2_ adrenergic receptors form dimers, whilst cannabinoid CB_1_ receptor and opsin form tetramers (*5*). The degree of oligomerisation is positively correlated with receptor density and with the presence of an agonist for some GPCRs, including the serotonin 5-HT_2c_ (*6*), dopamine D_2_ (*7*) and secretin receptors (*8*). The quaternary structures of GPCR oligomers were shown to be diverse, dynamic and modulated by ligand binding(*9, 10*). The biological importance of GPCR oligomerisation may lie in the possibility of functional communication between adjacent protomers (i.e. individual GPCR molecules) within these assemblies. Such communication is suggested e.g. by the multiphasic binding kinetics of an allosteric ligand to M2 receptors in native membranes, which was missing when the receptor was purified as monomers(*11*). Such intermolecular communications among GPCR protomers may be responsible for specific signalling pathways. Some recent super-resolution microscopy studies revealed that distinct clusters of GLP-1R (a Class B GPCR) form nanodomains to initiate endocytosis (*12*), and that G protein coupling and β-arrestin mediated internalisation are associated with the formation of CXCR4 oligomers (*13*). A recent single-molecule study links formation of μ-opioid receptor dimers with specific agonists and their downstream signals (*14*).

The detection of G protein oligomers has pointed towards a scenario in which a signalling complex may consist of an oligomeric assembly of both GPCRs and G proteins. Single particle analyses showed that a heterotetramer of A_1_-A_2a_ receptors exists as a dimer of dimers which simultaneously couples to both a Gi and a Gs protein (*15*), and that addition of agonist resulted in a supramolecular complex of four receptors and four G proteins (*16*). Similar to GPCR oligomerisation, that of G proteins is dynamic and subject to modulation by such factors as palmitoylation (*17*) and the presence of ligands(*18*).

A variety of GPCR dimer interfaces have been revealed by crystal structures: *e.g.* the symmetric interfaces of TM1,2,8/TM1,2,8 in the β_2_ adrenergic (*19*) and μ-opioid receptors (*20, 21*), and of TM5,6/TM5,6 as in the CXCR4 (*22*) and P2Y_12_ receptors (*23*). Asymmetric dimer interfaces have also been observed, e.g. TM1-3/TM5,6 in A_2a_ (*24*), and TM4,5/TM7 in the P2Y_1_ receptors (*25*). Lipids, *e.g.* cholesterol molecules and some fatty acid chains, are frequently seen at such dimer interfaces in *e.g.* P1Y_12_, A2aR and β_2_AR and others (*26*). Indeed, cholesterol was proposed to have an impact on GPCR oligomerisation either via alteration of bilayer properties, *e.g.* thickness, fluidity, acyl tail order, and/or by modulating the potential transmembrane interactions surface of the receptors by binding at specific lipid sites (*27*).

Molecular dynamics (MD) simulations, in particular coarse-grained (CG) simulations, provide a useful tool to investigate protein-protein interfaces and the impact of membrane lipids on GPCR oligomerisation. Rhodopsin was found to form interfaces with different association affinities corresponding to TM5/TM5 and TM1,2,8/TM1,2,8 interactions in CG simulations in POPC membranes (*28*). Addition of cholesterol molecules to pure phospholipid membranes shifted the dominant dimer interface of CXCR4 from TM1/TM5-7 to TM3,4/TM3,4 in CG simulations (*29*). Increasing the concentration of cholesterol in POPC membranes correlates with enhanced plasticity and flexibility of dimeric association of the serotonin_1A_ receptor, a modulatory effect exerted via both direct cholesterol binding and indirect membrane effects in CG simulations (*30*). Kinetics of the homodimerisation of opioid receptors have also been explored via CG simulations in a POPC/CHOL membrane and Markov state models, revealing that dimerisation of μOR is dynamic with kinetically distinct dimeric assemblies (*31, 32*).

Despite these efforts, our current understanding of GPCR oligomerisation still remains limited, reflecting a sparsity of structural information. Without a full characterisation of the quaternary structures of GPCR oligomers, it is difficult to understand how GPCR oligomerisation responds to receptor activation and signalling. Here, we provide a comprehensive characterisation of oligomerisation, quaternary structures, and kinetics of the Adenosine 2a receptor (A2a, a prototypical Class A GPCR) via extensive MD simulation data (~2.6 ms of CG-MD in total) using *in vivo* mimetic membranes. Our simulations reveal that both oligomer quaternary structures and the kinetics of A2aR oligomerisation are subject to modulations by the receptor‘s conformational state and by the lipid interactions of the receptor. Such responsiveness of A2aR oligomerisation suggests a combinatory allosteric modulation of GPCR signalling, in which the receptor may respond to the surrounding membrane environment to generate a unique oligomerisation profile favouring the formation of specific supramolecular complexes. This helps us to understand the structural details of GPCR signalling complexity at a larger scale, presenting new possibilities to manipulate GPCR function.

## Results

### MD simulations sample both association and dissociation of A2aR

We based initial simulations on large-scale membrane systems using a mixture of lipids to form an *in vivo* mimetic model of the plasma membrane environment within which multiple copies of the receptor can freely move, enabling both association and dissociation events to occur. We have focused on oligomerisation of the Adenosine A2a receptor (A2aR) as this receptor has been demonstrated experimentally to form dimers and higher order oligomers (*33, 34*). Thus we placed 9 copies (positioned and oriented randomly in membrane plane relative to one another) of the A2aR in a 45 × 45 nm^2^ membrane with 10 lipid species present (Fig. 1A) to simulate the oligomerisation process (i.e. the 9-copy systems). To study how the oligomerisation may change in response to receptor activation, we generated 3 ensembles of simulations for the 9-copy systems, one each for the receptor in the inactive state (PDB id 3EML), the active state (PDB id 5G53 receptor only) and the active state in complex with a mini Gs protein (PDB id 5G53 receptor and mini Gs). The simulation systems were modelled using the MARTINI coarse-grained force field (*35, 36*), including an elastic network to retain the receptor in a given conformation throughout the simulation, thus de-coupling the oligomerisation process from receptor conformational changes. We also performed simulations of the three conformational states with a higher protein density (~4% by area) by including of 16 copies of the receptor in a membrane of the same dimensions (termed 16-copy), as a number of studies (*e.g.* (*5*)) have shown that altering receptor density can alter GPCR oligomerisation. Given recent studies, both computational and experimental (*37–39*), demonstrating the interaction of phosphatidylinositol 4,5-bisphosphate (PIP_2_) with Class A GPCRs, we also performed simulations with setups identical to the 9-copy system but with a lipid bilayer devoid of PIP_2_ (9-copy NoPIP_2_) to see whether binding of PIP_2_ could modulate GPCRs via altering oligomerisation.

**Figure 1.**
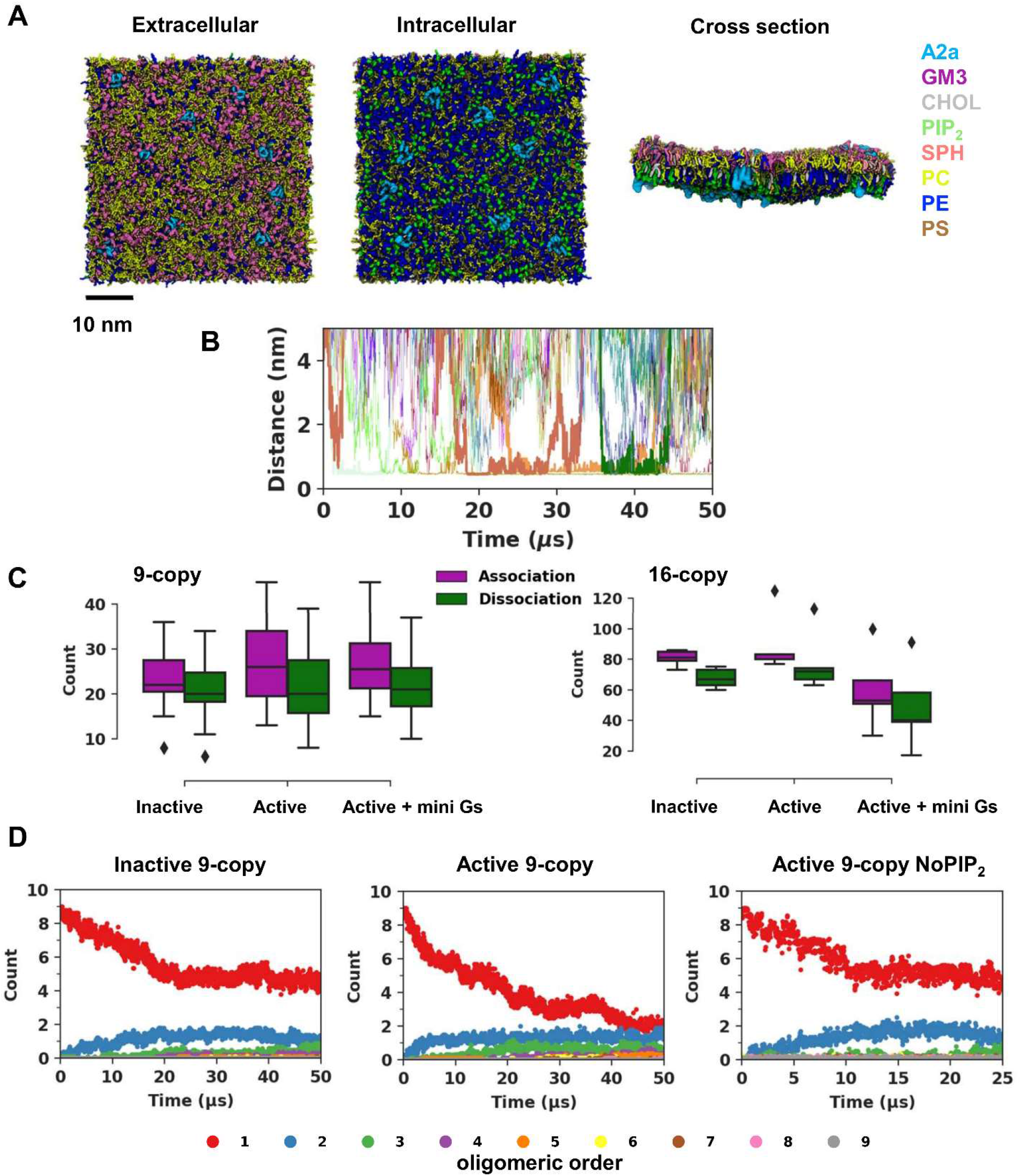
A2aR oligomerisation sampled by MD simulations using complex membranes. **(A)** System setup of A2aR oligomerisation simulations. The selected number of receptor molecules (pale blue) were randomly inserted into a mixed lipid membrane of area 45 × 45 nm^2^. Views of the systems from the extracellular and intracellular surfaces and in cross section are shown, with the lipid species present colour coded. Details of the different simulation systems setups are listed in Table 1. **(B)** The time evolution of minimum distance between pairs of receptors (from a simulations trajectory in the inactive state 9-copy simulation ensemble), illustrating that both association and dissociation events occur within the timescale simulated (selected events are highlighted by bold traces). **(C)** The number of association (defined as the smoothed minimum distance of a pair of proteins coming closer than 0.75 nm) and dissociation (defined as the smoothed distance of a pair of proteins separating to further than 0.75 nm) events sampled in each trajectory for each of the 9-copy and 16-copy system ensembles. **(D)** The associations and dissociations have led to a dynamic equilibrium as illustrated by the time course of oligomer formation for the inactive state 9-copy, active state 9-copy, and active state 9-copy NoPIP_2_ simulations. Data averages from all trajectories in the same simulation system were plotted to illustrate the ensemble trend. See SI Fig S2 for a full list.

Overall, 60 MD trajectories with a cumulative simulation time of more than 2.6 ms were generated (Table 1). The time evolution of the minimum distance between pairs of receptors demonstrated that multiple association and dissociation events occur within the timescales simulated, and protein-protein interactions of different durations were sampled (Fig. 1B and SI Fig. S1). On average, each trajectory sampled over 20 association or dissociation events in the 9-copy systems and over 60 in the 16-copy systems (Fig. 1C). The total number of association events in the 16-copy systems was approximately (16/9)^2^ times higher than in the 9-copy system, indicating a second order reaction. The ensemble data of the number of monomers, dimers, and higher order oligomers suggested that all nine simulation systems approached an equilibrium towards the end of simulations, the characteristics of which were dependent on the membrane environment and the receptor conformational state (Fig. 1D and SI Fig. S2). For example, comparing the 9-copy simulations of the Inactive vs. Active states, it can be seen that after 25 to 30 μs, the number of monomers drops to ~5 for the Inactive state compared to 2 or 3 for the Active state. The effect of omitting PIP_2_ from the simulation of the Active state of the receptor decreases the tendency of the receptor to oligomerise such that the number of monomers remained at ~5.

**Table 1.**
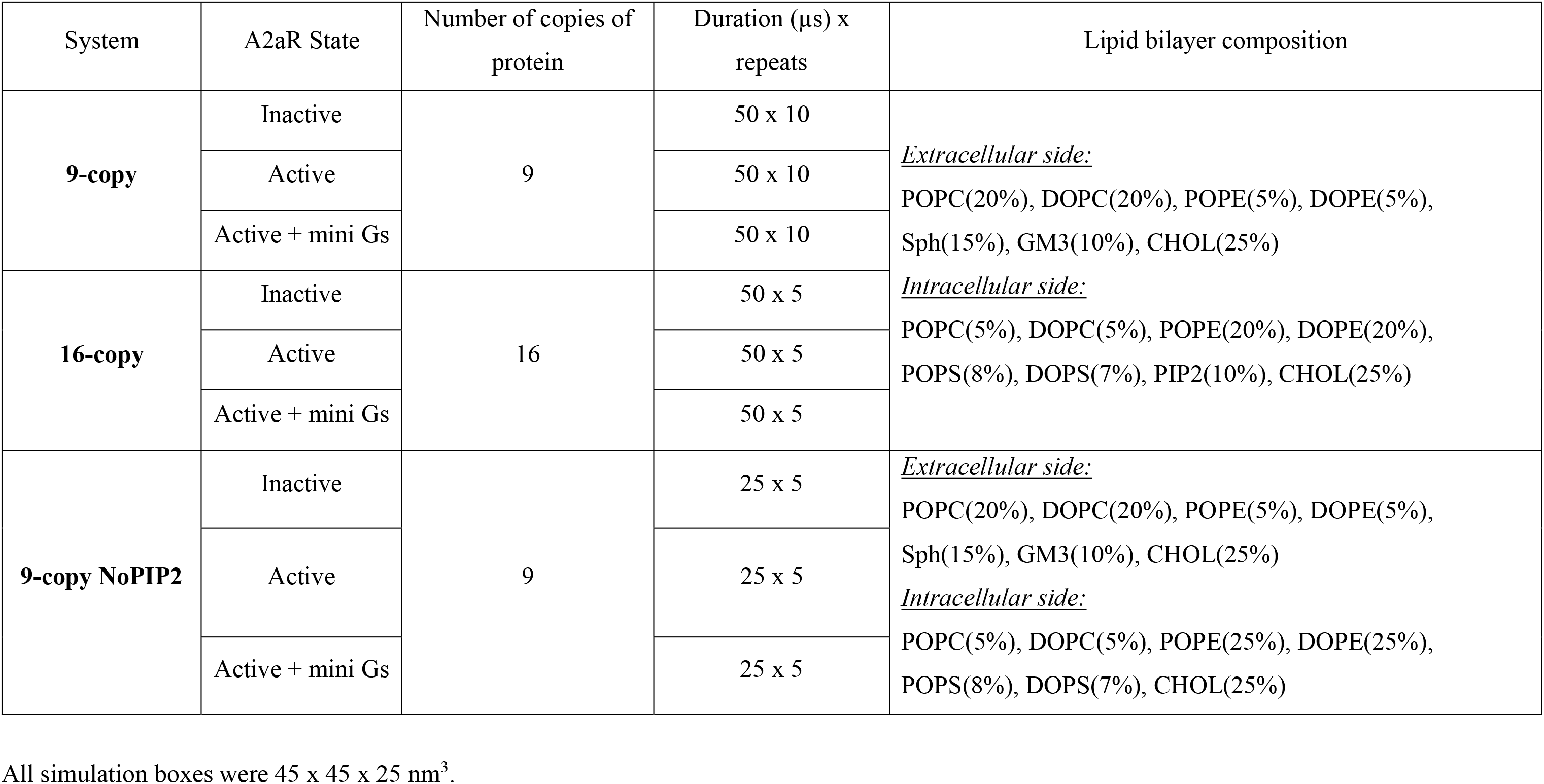
Overview of Simulations Performed

### Oligomerisation profiles depend on the conformational state of the receptor

Determining the distribution of oligomers within each simulation system showed that the active state favours oligomerisation (note: we use ‘oligomerisation’ to refer to both dimerisation and higher order oligomerisation) compared to the inactive state. Thus, for the active state, about 66% of the receptors were oligomeric compared to 40% for the inactive state in the 9-copy systems (Fig. 2). Not surprisingly, comparison of the 9-copy and 16-copy simulations indicates that a higher receptor density in the membrane favours oligomerisation. The presence of the mini-Gs protein bound to the receptor also shifted oligomerisation profiles in favour of higher order oligomers. Increased frequencies of higher order oligomers were seen for the mini Gs-coupled state in all three sets of simulations. The correlation of oligomerisation with both receptor activation and protein density is in agreement with observations on a number of GPCRs(*6–8*).

**Figure 2.**
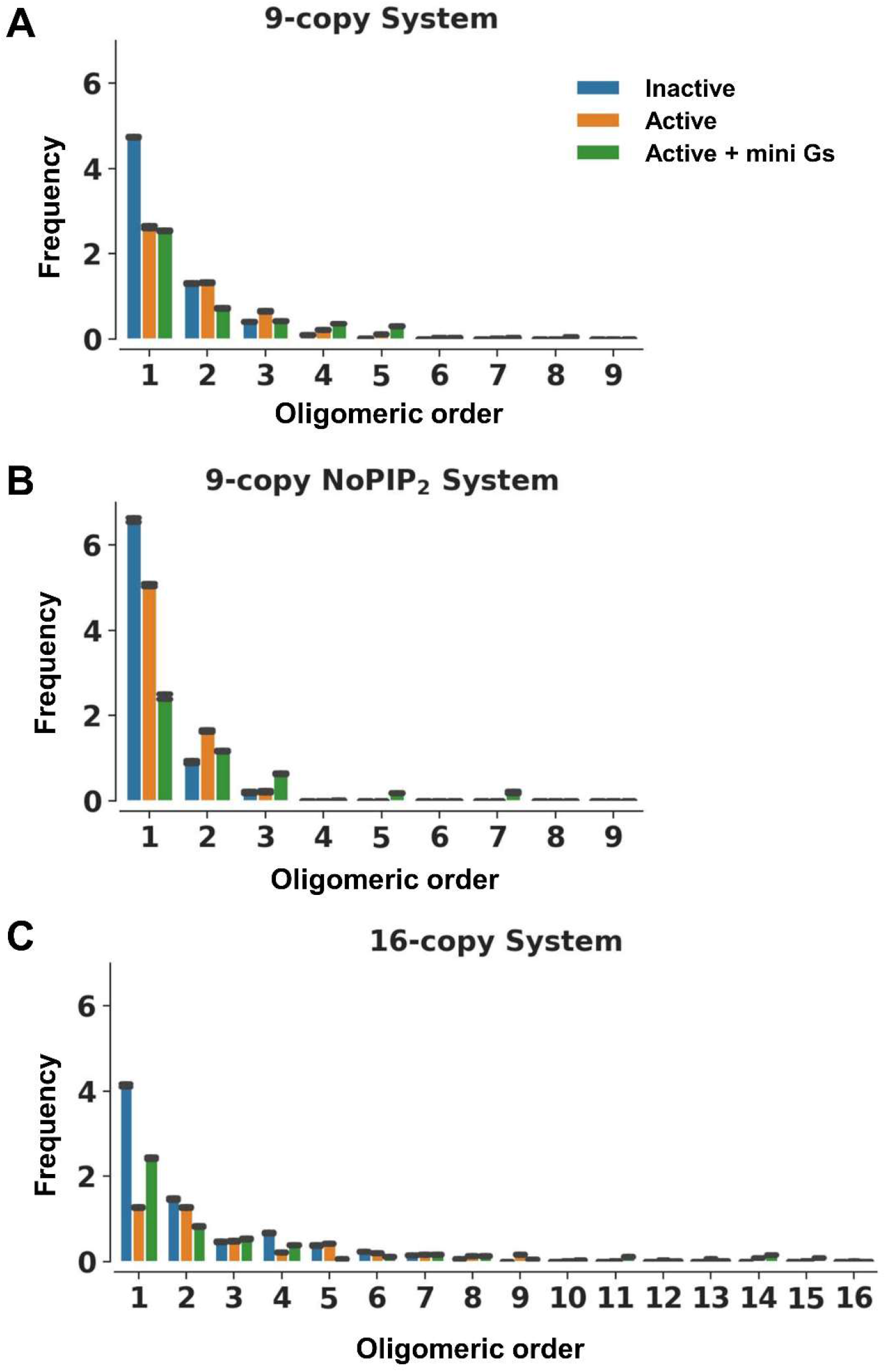
Oligomer distribution. The oligomer distributions are shown for the final 20 μs in the 9-copy simulation **(A)** and 16-copy simulations **(B)** and for the final 10 μs in the 9-copy NoPIP_2_ simulation **(C)**, estimated by counting the number of oligomers in the systems. The error bars (black) denote standard deviations along the trajectory time course.

#### Metrics

To better understand the oligomerisation landscape of A2aR, we used two metrics to describe their oligomerisation profile: (i) the population of various oligomeric configurations (i.e. oligomer quaternary structures) to measure their relative likelihood, and (ii) the residence times of these configurations to provide a measure of their relative stability (and thus the relative strength of the corresponding protomer interactions). We therefore clustered all the oligomer structures using a technique that is invariant to permutations of molecular indexing and calculated the residence time of each oligomer cluster based on their survival functions (see Fig 3A and Methods).

**Figure 3.**
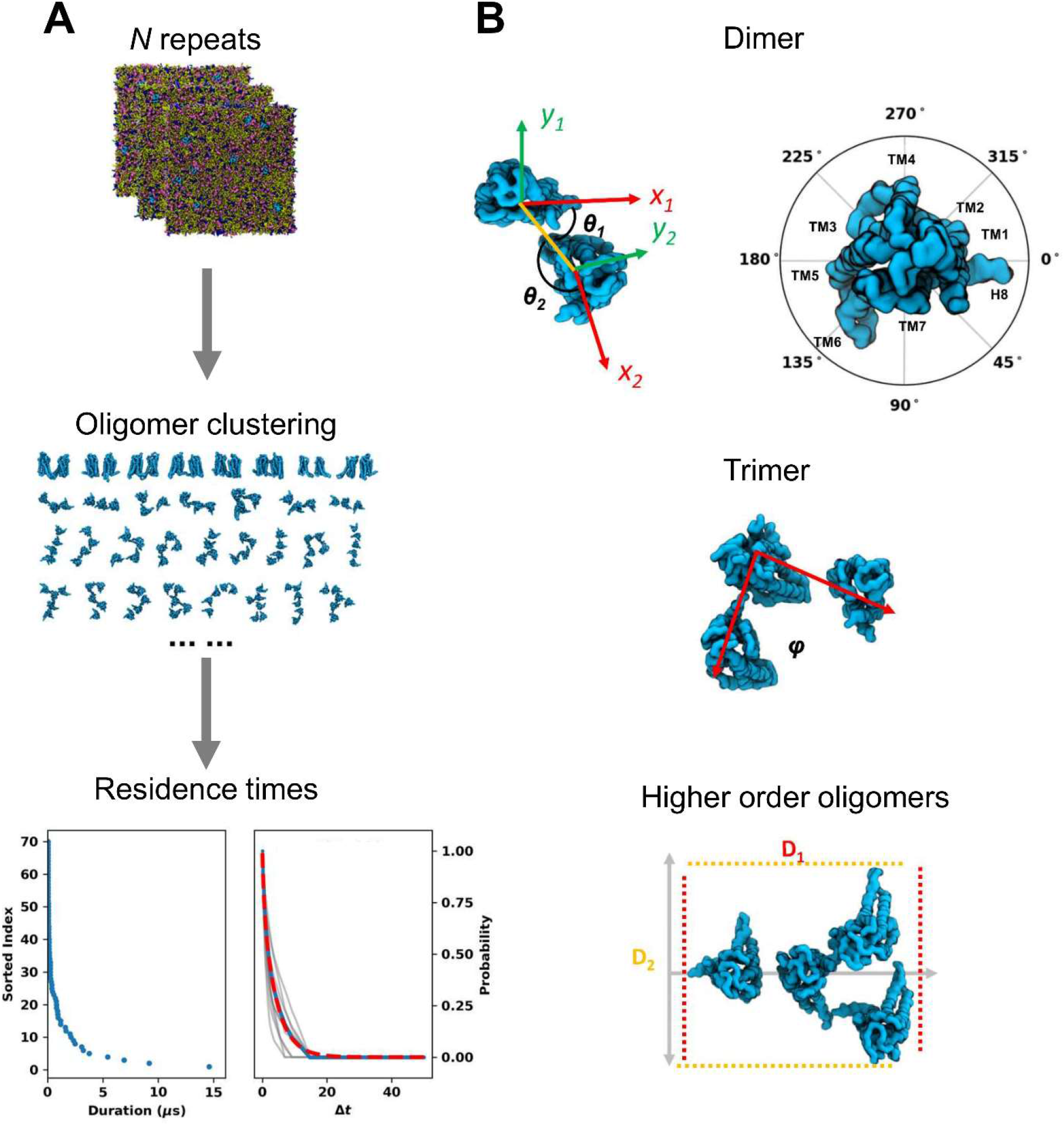
Characterisation of oligomer quaternary structures. **(A)** The oligomer quaternary structures from the same simulation set were clustered to identify the various oligomeric configurations. For the calculation of oligomer residence time, durations of each configuration were collected and sorted (blue dots in the left panel). A normalised survival function as a function of Δ*t* (blue dots in the right panel) were modelled based on the sorted durations. A biexponential (red dashed line in the right panel) was used to fit to the survival function to obtain *k*_*off*_. To estimate the confidence of the calculated *k*_*off*_ values, survival functions based on bootstrapped durations were modelled (grey lines in the right panel) from which standard deviations were calculated. **(B)** The following metrics were used to describe the oligomer configurations: for dimers, two binding angles (*θ*_*1*_, *θ*_*2*_) such that each of the angles describes the relative position of the dimer interface to the principal axis of that monomer that is parallel to H8 in a clockwise direction; for trimers, the bending angle *φ* defined by the centre of mass of the three monomers; and for tetramers and pentamers, the projected lengths *D*_*1*_ and *D*_*2*_ on their 1^st^ and 2^nd^ principal axes.

To assist the calculation of populations of the different oligomeric configurations, we used the following metrics to describe these quaternary structures (Fig. 3B): (i) for dimers, the binding angles (θ_1_, θ_2_)(*40*), which describe the angle defined by the dimer interface and the principal axis of the monomers that is parallel to H8; (ii) for trimers, a bending angle *φ* defined by the centres of mass of the three monomers; and (iii) for tetramers and pentamers, the projected lengths D_1_ and D_2_ onto their 1^st^ and 2^nd^ principal axes respectively. For the 9-copy systems, oligomers of orders lower than pentamers corresponds to 99.97%, 99.93% and 91.68% of the total populations of the inactive, active, and active + mini Gs states respectively.

#### Dimers

Comparison of the dimeric populations in the 9-copy systems revealed that A2aR dimerisation is sensitive to the conformational state of the receptor. Thus inactive state dimers predominantly showed interfaces: (TM1,H8//TM1,H8) (20°, 20°); (TM3,TM5//TM7,H8) (40°, 170°); and (TM3,TM5,ICL2//TM3,TM5,ICL2) (200°, 200°) (where ICL = intracellular loop, TM = transmembrane helix and A//B indicates an interface between surface A and surface B). When the receptor was in the active state, dimerisation around (20°, 20°) and (40°, 170°) became less frequent whereas the dominant interfaces shifted to those involving ICL3, i.e. interfaces around (TM3,TM5,ICL3//TM7, H8) (70°, 200°) and (TM5,TM6,ICL3//TM5,TM6,ICL3) (130°, 130°) (Fig. 4A).

**Figure 4.**
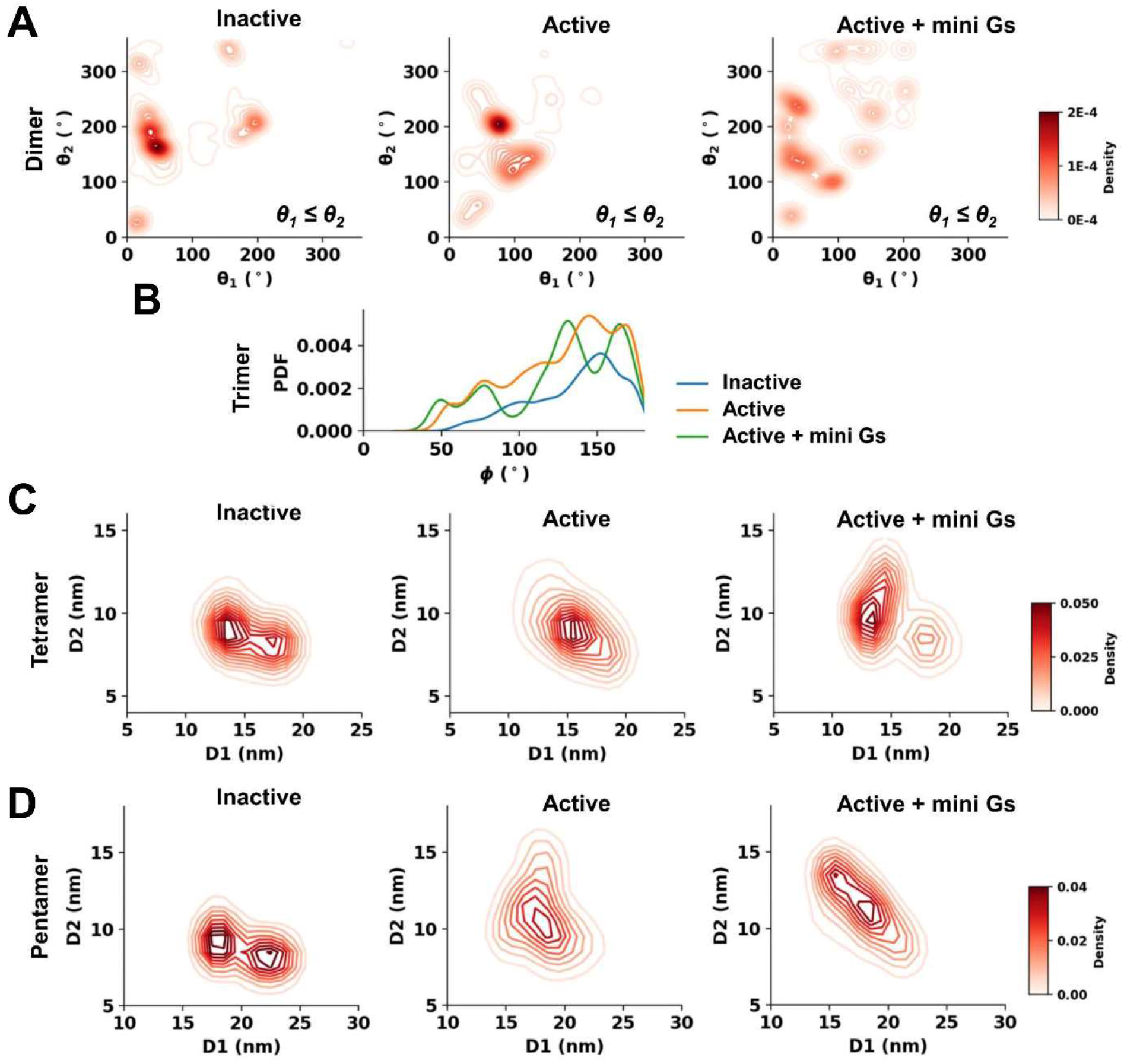
Population profiles of oligomeric configurations in the 9-copy ensemble. Distributions of oligomeric configuration metrics defined in Fig. 3B are shown in **(A)** for dimers, **(B)** for trimers, **(C)** for tetramers and **(D)** for pentamers.

From the dimer residence times, the most stable dimer interface in the inactive state was TM1,H8//TM5,ICL3 (190°, 335°) with a residence time of 14 μs but only an intermediate population (Fig. 4A, 5A and SI Table S1). This dimer interface is observed in an inactive state of the A2aR crystal structure (PDB id 4EIY). This suggests that residence time, as an indication of oligomer stability, may be a better metric than relative abundance of an oligomeric configuration to predict dimer interfaces in crystallography from simulations. Indeed, the dimer interfaces seen in crystal structures of a couple of other Class A GPCRs also ranked highly in terms of residence times for the inactive state. For example, TM3,TM5,ICL2//TM3,TM5,ICL2 (186°, 196°) with a residence time of 10 μs is seen in the crystal structure of CXCR4 (PDB id 3ODU), and TM1,H8//TM1,H8 (30°, 30°) with a residence time of 6 μs is seen in inactive state structures of the β_2_Ad (PDB id 2RH1) and the μ-opioid receptors (PDB id 5C1M; SI Table S1).

**Figure 5.**
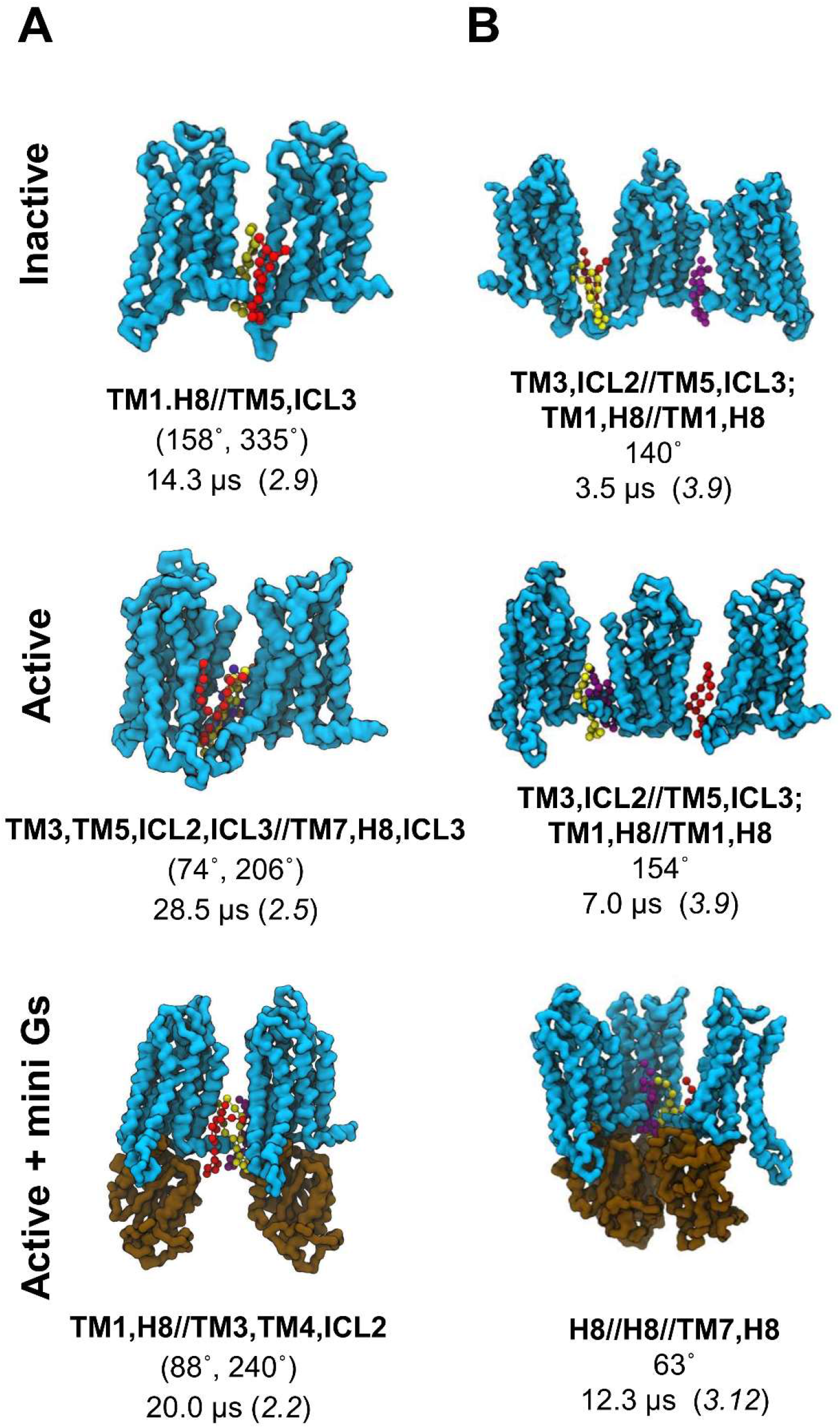
Oligomeric configurations with the longest residence times in the 9-copy ensembles. The dimeric **(A)** and trimeric **(B)** structures with the longest residence times in each conformational state are shown, with the backbone beads of the receptor in cyan surface and the mini Gs in brown. PIP_2_ molecules at the association interfaces are shown in ball and sticks with different colours. Below each of structure are the association interfaces noted in bold (where ICL = intracellular loop, TM = transmembrane helix and A//B indicates an interface between surface A and surface B), along with the average values of the descriptive metrics of oligomer configurations (see text and Fig. 3B for details), and the oligomer residence time which is followed by the cluster id label in brackets. Also see SI Fig. S3-S4 and SI Table S1.

Stabilities of dimer interfaces in the active state were enhanced (Fig. 6C), with a mean residence times of 6.2 μs (range 1-28 μs), compared to 5.3 μs (range 0.8-14 μs) for the inactive state dimers. The most stable dimer in the active state with the interface TM3,TM5,ICL2,ICL3//TM7,H8,ICL3 (74°, 206°) had a residence time of 28 μs (Fig. 5A). This resembles a stable inactive interface TM3,TM5//TM7,H8 (41°, 172°) with a residence time of 8 μs (SI Figure S3) but contained additional inter-protomer interactions at the intracellular side of TM6 and ICL3 relative to its inactive counterpart. This stable active interface also had the largest relative population. Another active highly populated dimer interface, TM5,TM6,ICL3//TM5,TM6,ICL3 (120°, 153°), ranked second in terms of residence times (10 μs). A similar dimer interface can be found in the inactive state at TM3,TM5//TM5,ICL3 (96°, 164°) in which inter-protomer contacts of the intracellular side of TM6 in the active state are replaced by TM5 in the inactive state, reducing the residence time to 2 μs. The active state dimer interface ranked the 3rd in residence time (9 μs; TM1,H8//TM1,H8 (35°, 46°); SI Fig. S3) again showed increased stability relative to its inactive counterpart (6 μs).

**Figure 6.**
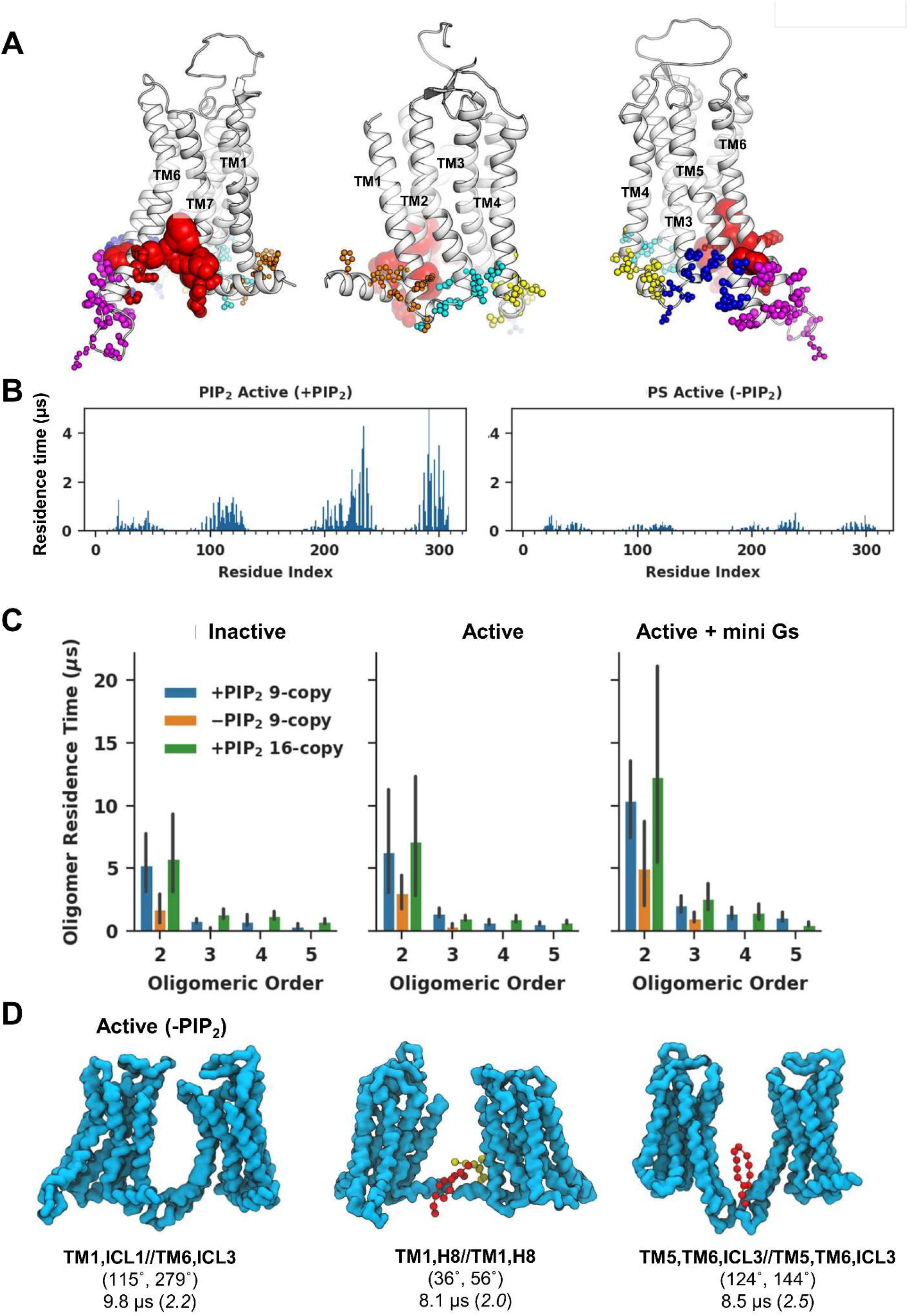
The influence of PIP_2_ on A2aR oligomerisation. **(A)** PIP_2_ binding sites on the active state A2aR. The six binding sites calculated using PyLipID (https://github.com/wlsong/PyLipID) are displayed using different colours. Residues in each binding site are shown as spheres with radii proportional to their PIP_2_ residence times for the active state 9-copy simulations. **(B)** Comparison of (left panel) PIP_2_ residence times vs. residue number in active state PIP_2_-containing simulations (9-copy) with (right panel) PS residence times in the active state PIP_2_-free simulations (9-copy NoPIP_2_). **(C)** Comparison of the average residence time as a function of oligomeric order between the three membrane environments (PIP_2_-containing, 9-copy; PIP_2_-free, 9-copy; and PIP_2_-containing, 16 copy) for the 3 states of the A2aR. Bar heights denote the average residence time of the oligomeric configurations from the same oligomeric order and the error bars show the 95% interval of 1000 bootstrapped samples. **(D)** Active state dimer configurations with the 3 longest residence time from PIP_2_-free simulations. The backbone beads of the receptor are shown in cyan. The PS molecules at the association interfaces are shown in ball and sticks with different colours. Below each of the structures are the association interfaces in bold, the average values of the binding angles, and the oligomer residence time which is followed by the cluster id label in brackets.

Binding of the active receptor to the mini Gs strengthened the dimer association as revealed by increased hot spots in the population distribution (Fig 4A). The average residence time of dimers was correspondingly increased to 10 μs (range 3-20 μs; Fig 6C). The enhanced stability resulted from the interactions between the receptor of one protomer and mini Gs of the other in addition to interactions between two receptors. The dimer association hot spots in the mini Gs-coupled state merged those from both the inactive and active states, suggesting that the dimer associations were more promiscuous, however, the stabilities of dimer interfaces, i.e. the ranking of residence times, agreed reasonably well with those in the active state (SI Table S1), indicating that the association stability was, to a large extent, governed by the activate state of the receptor.

#### Trimers and higher order oligomers

The residence times of trimers and higher order oligomers were much lower than for dimers, indicating that the oligomerisation at higher orders was dynamically unstable (Fig 6C). These oligomers presented more compact quaternary structures in the active state compared to the inactive state, i.e. more bent trimers with smaller φ or more branched/closed higher order oligomers with similar values of D1 and D2 (Fig 4B-4D, SI Fig S4). This shift resulted from the opening at the intracellular side of TM6 and TM5 in the active state that increased the area of the intracellular receptor surface formed by H8-TM6-ICL3 and ICL2-TM5-ICL3. Such a shape made branched/closed quaternary structures more energetically favourable. Dynamic and compact oligomeric structures have been reported for the Ste2 receptor by FRET (*9*). The coupling of mini Gs further enhanced this shift to more compact configurations in which some of the signalling partners made contacts with one another. Supramolecular organisation of oligomeric GPCRs in complex with oligomeric G proteins have been reported based on single-particle photobleaching experiments(*16*). The interactions between signalling partners in such supramolecular assemblies were suggested to enable communications between them and hence may provide a structural explanation to the allosteric modulations in GPCR oligomers (*16*).

### PIP_2_ enhances protein-protein associations and changes association interfaces

Inspection of the 9-copy simulation trajectories revealed that the protein-protein associations were mediated by lipid molecules, predominantly by PIP_2_ molecules as shown in Fig 5. The radial distribution of lipids around the receptor (see SI Fig. S5) confirmed that PIP_2_ accumulated around the receptor. We used graph theory and community analysis (see Methods for details) to characterise the binding sites and binding kinetics. Six PIP_2_ binding sites on A2aR were pinpointed based on the interactions of lipid headgroup beads (Fig 6A) and the lipid residence times at these binding sites were calculated (Table 2). The strengths of PIP_2_ interactions with A2aR (as measured by residence times) correlated well with the stabilities of dimeric associations they mediated. Thus, for the inactive state, three strong (i.e. long residence time) PIP_2_ binding sites, TM1_H8, TM3_TM5 and TM6_TM7, contributed to the most stable dimer interfaces in this state, i.e. (TM1,H8//TM1,H8) (20°, 20°); (TM3,TM5//TM7,H8) (40°, 170°); and (TM3,TM5,ICL2//TM3,TM5,ICL2) (200°, 200°) (see above). For the active state, the greatest increase in PIP_2_ residence time was seen at site TM6_TM7, which went from 2.4 μs in the inactive state to 12 μs in the active state. Similar increase was seen in the PMF calculation of PIP_2_ binding at this TM6-TM7 site in simulations of a single copy of A2aR (*37*), suggesting that this increase of PIP_2_ stability is independent of A2aR oligomerisation. This PIP_2_ site mediated the most stable dimer interface (TM3,TM5,ICL2,ICL3//TM7,H8,ICL3) of the active state (Fig 5A). Two other PIP_2_ binding sites, TM5_TM6_ICL3 and TM3_TM5, also showed substantial increase of PIP_2_ residence times on going from inactive to active, again correlated with a shift of dimer associations to those involving ICL3 in the active state. In the mini Gs-coupled state, PIP_2_ residence times at all binding sites increased significantly, which again agreed well with the increase of protein-protein association residence times in this state (Table 2 and Fig 6C). The overlapping nature of the PIP_2_ binding sites and the protein/protein interfaces, and the strong correlation of PIP_2_ and protein/protein residence times suggests that PIP_2_ interactions are a key determining factor in driving A2aR oligomerisation as observed in the simulations.

**Table 2.** Anionic Lipid Binding Sites in 9-Copy Simulations

| Binding site<sup>§</sup> | TM1_H8 |  |  | ICL1_TM4 |  |  | TM3_TM4_ICL2 |  |  | TM3_TM5 |  |  | TM5_TM6_ICL3 |  |  | TM6_TM7 |  |  |
| --- | --- | --- | --- | --- | --- | --- | --- | --- | --- | --- | --- | --- | --- | --- | --- | --- | --- | --- |
| A2aR state* | I | A | AmG | I | A | AmG | I | A | AmG | I | A | AmG | I | A | AmG | I | A | AmG |
| **PIP<sub>2</sub> binding sites in PIP<sub>2</sub>-containing simulations** |  |  |  |  |  |  |  |  |  |  |  |  |  |  |  |  |  |  |
| **Residence time (μs)** | 1.4 | 1.6 | 20.2 | 0.6 | 1.0 | 21.3 | 0.5 | 0.5 | 27.1 | 1.5 | 3.2 | 20.4 | 0.8 | 4.9 | 8.1 | 2.4 | 12.1 | 31.3 |
| **Number of PIP<sub>2</sub> molecules** | 4.4 | 2.9 | 5.4 | 2.7 | 2.1 | 3.1 | 3.0 | 3.0 | 3.3 | 3.8 | 3.6 | 5.1 | 4.2 | 5.6 | 5.1 | 3.9 | 4.1 | 3.4 |
| **DOPS binding sites in PIP<sub>2</sub>-free simulations** |  |  |  |  |  |  |  |  |  |  |  |  |  |  |  |  |  |  |
| **Residence time (μs)** | 0.2 | 0.7 | 1.7 | 0.2 | 0.17 | 1.2 | 0.2 | 0.2 | 3.5 | 0.06 | 0.09 | 2.9 | 0.2 | 0.2 | 2.4 | 0.2 | 0.5 | 3.0 |
| **Number of DOPS molecules** | 1.6 | 1.6 | 1.6 | 1.3 | 1.4 | 1.6 | 1.6 | 1.6 | 1.7 | 1.5 | 1.5 | 1.6 | 1.6 | 1.6 | 1.4 | 1.4 | 1.8 | 1.7 |

<sup>§</sup> The binding sites are illustrated in Fig. 6B, to which the colour coding used above corresponds. Binding sites were calculated based on the interactions of headgroup beads.
\* I = inactive; A = active; AmG = active + mini Gs.

#### NoPIP2 simulations

To further explore the influence of PIP_2_ interaction on A2aR oligomerisation, we calculated the oligomerisation residence times and lipid interactions in the 9-copy NoPIP_2_ simulations (i.e. the simulation in a bilayer omitting PIP_2_). Regardless of the conformational state of the receptor, the stability of protein-protein associations was significantly reduced for all oligomeric orders (Fig. 6C). This reduced stability had led to a lower overall degree of oligomerisation compared to the PIP_2_-containing systems (Fig 2). Tetramers or larger oligomers were rarely seen in these simulations. Lipid radial distributions for the NoPIP2 simulations (SI Fig. S5) showed a second peak from PS, with a lower height than for PIP_2_ in the PIP_2_-containing simulations. Visual inspection of simulations confirmed that PS molecules were often seen at the protein/protein interfaces (Fig. 6D). The distribution of PS interactions with the A2aR was similar to that of PIP_2_ but with much lower residence times, especially at TM6 and TM7 (residue index 220-240 and 280-310; Fig 6B). Six PS binding sites, similar to those of PIP_2_, were identified, with faster lipid dissociation kinetics than PIP_2_. The PS binding site at TM6_TM7 did not show a prominent increase of lipid residence time in the active state comparing to the inactive state, in contrast with the large increase of PIP_2_ residence time at that site (see above). Accordingly, the strongest dimer interface (TM3,TM5,ICL2,ICL3//TM7,H8,ICL3 (74°, 206°)) in the active PIP_2_-containing simulations was rarely sampled in the active NoPIP_2_ simulations (SI Fig S6). The binding site at TM3_TM5 exhibited the weakest PS interactions among the 6 binding sites in both inactive and active states of the NoPIP_2_ simulations but showed one of the strongest PIP_2_ interactions (Table 2). This difference correlated with the absence of the stable dimer interface TM3,TM5,ICL2//TM3,TM5,ICL2 (186°, 196°) in the NoPIP_2_ simulations. Taken together, our data show that the PIP_2_ molecules can stabilise A2aR protein-protein associations, and lead to specific association interfaces that are mediated by strong PIP_2_ interactions with the receptor surface.

### Markov state models reveal more dynamic oligomerisation network in the active state and in the presence of PIP_2_

To explore the kinetics of A2aR oligomerisation in more detail, we constructed Markov state models (MSMs). MSMs can be used to model oligomerisation in terms of a network of transitions between oligomeric states, based on information extracted from MD simulations (*41*). We used the oligomeric composition of the MD simulation systems, i.e. the number of monomers, dimers and trimers etc. in the system, for MSM state decomposition to provide a physical description of the system-states. An MSM was estimated for each set of simulations using PyEMMA (*42*), which reweights the transitions such that the equilibrium kinetics and stationary distributions can be recovered. After statistical validation, the MSMs were used to compute equilibrium probabilities, kinetics, and oligomerization networks (see Methods for details).

The MSMs thus constructed revealed that the inactive state 9-copy systems preferably stayed at the following system-states: *1*^*9*^ (i.e. 9 monomers), *1*^*7*^-*2*^*1*^ (i.e. 7 monomers and 1 dimer) or *1*^*3*^-*2*^*3*^ (i.e. 3 monomers and 3 dimers), with lifetimes over 8 μs. In contrast, for the active state 9-copy systems the predominant system-states were *1*^*2*^-*2*^*2*^-*3*^*1*^, *1*^*1*^-*2*^*1*^-*3*^*2*^, *1*^*2*^-*2*^*1*^-*5*^*1*^ or *1*^*1*^-*3*^*1*^-*5*^*1*^, with lifetimes over 8 μs. The active state receptor also saw an increase in lifetime and population of system-states containing higher order oligomers. Accordingly, the lifetime of the monomeric state (*1*^*9*^) decreased to 6 μs in the active state (SI Table S3). The mini Gs-couple 9-copy systems showed extended lifetimes of 14 μs of *1*^*2*^-*3*^*1*^-*4*^*1*^, *1*^*4*^-*5*^*1*^, and had 5 system-states containing oligomers larger than trimers with lifetimes over 8 μs. The long lifetimes of these two MSM system-states suggests that the corresponding oligomeric configurations were favoured by the presence of the mini Gs either via forming stable interactions between the receptor and mini Gs or via forming stable interactions among the mini Gs proteins. The lifetime of the fully monomeric system-state was further decreased to 5 μs in this mini Gs-coupled state. Reaction rates revealed a more dynamic oligomerisation network for the active state receptor, with faster transitions between states (Fig 7). In contrast, for the inactive state the rates of transition between system-states were much smaller with the exception of the transition from *1*^*1*^-*2*^*1*^-*6*^*1*^ to *1*^*4*^-*2*^*1*^-*3*^*1*^. Simulated MSM trajectories using a Monte Carlo algorithm and MSM transition probabilities revealed that the system with active A2aR has more heterogeneous populations of oligomers with more dynamic transitions among higher order oligomers compared to that with the inactive receptors (SI Figure S7).

**Figure 7.**
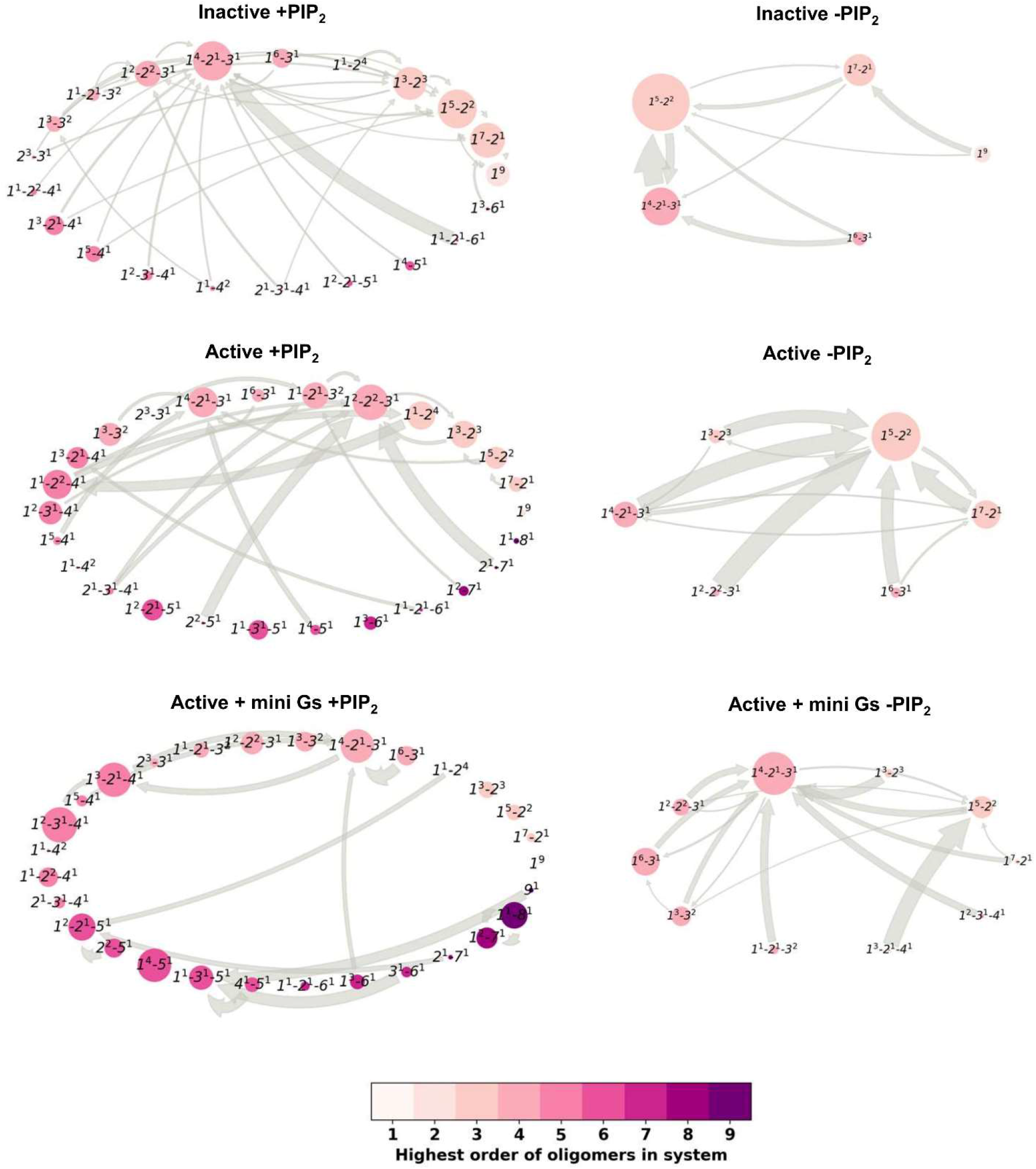
Markov state models of A2aR oligomerisation. The models calculated the kinetics of the oligomerisation by monitoring the evolution of A2aR oligomers in the membrane. The oligomerisation states are labelled as *A*^*a*^-*B*^*b*^-*C*^*c*^ in which *A, B, C* denote the oligomeric orders present in the membrane and *a, b, c* denote the number of oligomers of the corresponding order. The thickness of the arrows is proportional to the corresponding reaction rate (only reaction rates >50 s^−1^ are shown) and the size of the circles to the equilibrium distributions. Reaction rates were calculated as the reciprocal of the corresponding mean first passage time. To assist visualisation, the circles are coloured based on the highest oligomer order the representing state contains. MSM trajectories of 10 ms based on the MSM transition probabilities can be found in SI Figure S7.

The oligomerisation networks in the 9-copy NoPIP_2_ systems were much smaller and simpler due to the absence of larger oligomers (Fig 7). The simpler transition networks have led to more homogeneous oligomer populations comprised of monomers, dimers and trimers (SI Figure S7). The residence time of the inactive all-monomer system state was 11 μs, longer than its counterpart in the PIP_2_-containing simulations, indicating a low tendency for oligomerisation regardless of the presence of PIP_2_. Similar to the PIP_2_-containing simulations, enhanced oligomerisation, i.e. decreased residence time of monomeric system state (5 μs), was seen for the active receptors. In particular, dimers were seen in many of the oligomeric system states. Taken together, the MSMs revealed that both the presence of PIP_2_ in membranes and receptor activations have led to enhanced oligomerisation of A2aR with more dynamic transitions and heterogeneous oligomer populations.

## Discussion

We have provided a comprehensive characterisation of A2aR oligomerisation configurations and kinetics by large-scale unbiased CG simulations of systems containing multiple copies of the receptors on multi-microsecond timescales, and subsequent construction of Markov state models. One of the key findings from this study is that A2aR oligomerisation was enhanced by receptor activation. The conformational changes associated with receptor activation, especially the outward tilt of TM6, shifted the oligomerization interfaces to those involving ICL3. Our results are in agreement with the mutagenesis study that showed a critical role of ICL3 in formation of configurations related to the functions of GPCR oligomers(*43*). The enhanced oligomerisation by receptor activation was accompanied by a more dynamic oligomerisation network with more diverse oligomer populations. The diverse population of oligomers may serve as a pool of keys that unlock (bind to) the various signalling partners, either in monomeric or oligomeric forms. The specificity of this key-lock effect has been demonstrated by the recent Ste2 homodimer structure with a dimer interface of TM1,TM2,TM7//TM1,TM2,TM7 which allowed for binding of a Gpa1 dimer but would have created serious steric clashes for G protein dimers seen in mammalian GPCRs (*44*). Hence, changes to the profiles of the oligomer populations could lead to changes in their coupled partners and the signalling profiles of the receptor. Receptor activation also led to a more connected oligomerisation network, making oligomerisation in this state more responsive to changes in *e.g.* the membrane environment. Given that outward tilt of TM6 upon activation is a common feature in Class A GPCRs, we suggest that the enhanced oligomerisation might be an element of a shared mechanism for receptor activation.

The interactions of membrane lipids, *e.g.* cholesterols and polyunsaturated fatty acids, have been reported to affect GPCR oligomerization(*45, 46*), via altering the receptor surface and intercalating between protomers(*27*). Here we report a profound effect of PIP_2_, previously shown to interact with a number of Class A GPCRs (*37–39*), on A2aR oligomerisation. We observed a direct correlation between the stability of receptor association interfaces and the strength of PIP_2_ interaction at these locations, (*e.g.* the increase of stability of interface TM3,TM5,ICL2,ICL3//TM7,H8,ICL3 and the increase of PIP_2_ residence time at binding site TM6_TM7). The impact of PIP_2_ on A2aR oligomerisation was further confirmed by comparisons to the NoPIP_2_ simulations, in which receptor oligomer residence times were significantly reduced whilst the dominant PIP_2_-mediated interfaces were absent. Given that our previous simulation studies have shown that the interactions of PIP_2_ are largely conserved across Class A GPCRs(*38*), we suggest that the effect of PIP_2_ on GPCR oligomerisation may also be shared. Such lipid interactions could be important in determining GPCR oligomerisation *e.g.* in functionally important nanodomains (*47*) with a high local concentration of PIP_2_(*48*). It is also possible that the intrinsically disordered C-termini of GPCRs (*49*) may play a role in interactions with anionic lipids such as PIP_2_, especially as they contain multiple basic residues (*50*).

In the active + mini Gs systems, the compact oligomeric configurations suggest that communication between mini Gs or other signalling partners may be possible in these dynamic supramolecular assemblies. The recent Ste2 homodimer structure in complex with two G proteins showed exchange of dynamics between the two G proteins via interactions between the α-subunit in one protomer and the β-subunit in the other (*44*). Such ‘lateral’ communications could lead to different signalling outcomes and may be the structural basis for the diversified profiles of GPCR signalling. Indeed, many studies have suggested that the oligomerisation of GPCRs may initiate signalling pathways different from those activated by the corresponding monomers. For example, recent single-molecule imaging techniques have revealed that G proteins recruited by monomers and dimers of the β_2_AR elicited different signalling pathways (*51*); the block of recruitment of G protein to β_2_AR dimers by the inverse agonist isoproterenol suggested that the dimers in this case were responsible for the basal activity of β_2_AR; and the specific effect of FTY720-P on S1PR1 was demonstrated to be dependent on receptor oligomerisation causing a short-lived Gαi-mediated intracellular response that differed from that of the endogenous agonist S1P(*52*).

From a methodological perspective, simulation studies using the MARTINI force field have provided valuable insights into association of membrane proteins (*28, 32, 53–57*) and into protein-lipid interactions (*58–60*). However, the current MARTINI model has some limitations (*61, 62*) which may sometimes lead to overestimation of direct protein-protein interactions in the aqueous phase. Whilst the Inactive and Active simulation ensembles showed reasonable association-dissociation behaviour, in part due to the involvement of lipids in the protein-protein interface, the Active + mini Gs ensembles displayed a lower frequency of dissociation possibly resulting from direct interactions of mini Gs in the aqueous phase. This low sampling of dissociations increased the uncertainties for the properties calculated for the Active + mini Gs state. Future studies of GPCR oligomerisation using MARTINI 3 (http://cgmartini.nl/index.php/martini3beta) may provide further information on *e.g.* the impact of G protein coupling on GPCR oligomerisation.

## Conclusions

Reflecting upon our simulation results in the context of experimental studies (*51, 52, 63*), we suggest that GPCR oligomerisation may provide greater functional flexibility for the receptor signalling array via generating multiple dynamic supramolecular complexes that could initiate different signalling outcomes (Fig 8). From our simulations, we observed enhanced oligomerisation with more connected networks in the active state which resulted in an array of oligomeric configurations capable of coupling to various configurations of oligomeric mini Gs or multiple copies of monomeric mini Gs. In addition, the oligomerisation energy landscape was sensitive to the membrane environment. Given that the coupling of G proteins to GPCRs is highly efficient, involving direct collisions with no intermediate(*64, 65*), the array of functional supramolecular complexes of GPCRs complexed with G protein(s) would be dependent on the array of oligomeric configurations presented by the activated receptor. Our simulations have revealed an additional level of complexity of GPCR allosteric modulation by receptor oligomerisation, whereby the receptor can generate a specific array of oligomers according to the environment that would lead to specific signalling complexes and hence functional outcomes. This hypothesis provides a structural explanation for the observation of multiple pharmacological profiles of GPCRs (*66*), and potentially expands the druggable space to include the protein-protein association interface and allosteric sites corresponding to the protein-lipid interface (see the discussion in (*26*)). Understanding the mechanisms of combinatory modulation of GPCR oligomerisation and the role therein of lipid interactions, may present new opportunities for therapeutic targeting of GPCRs.

**Figure 8.**
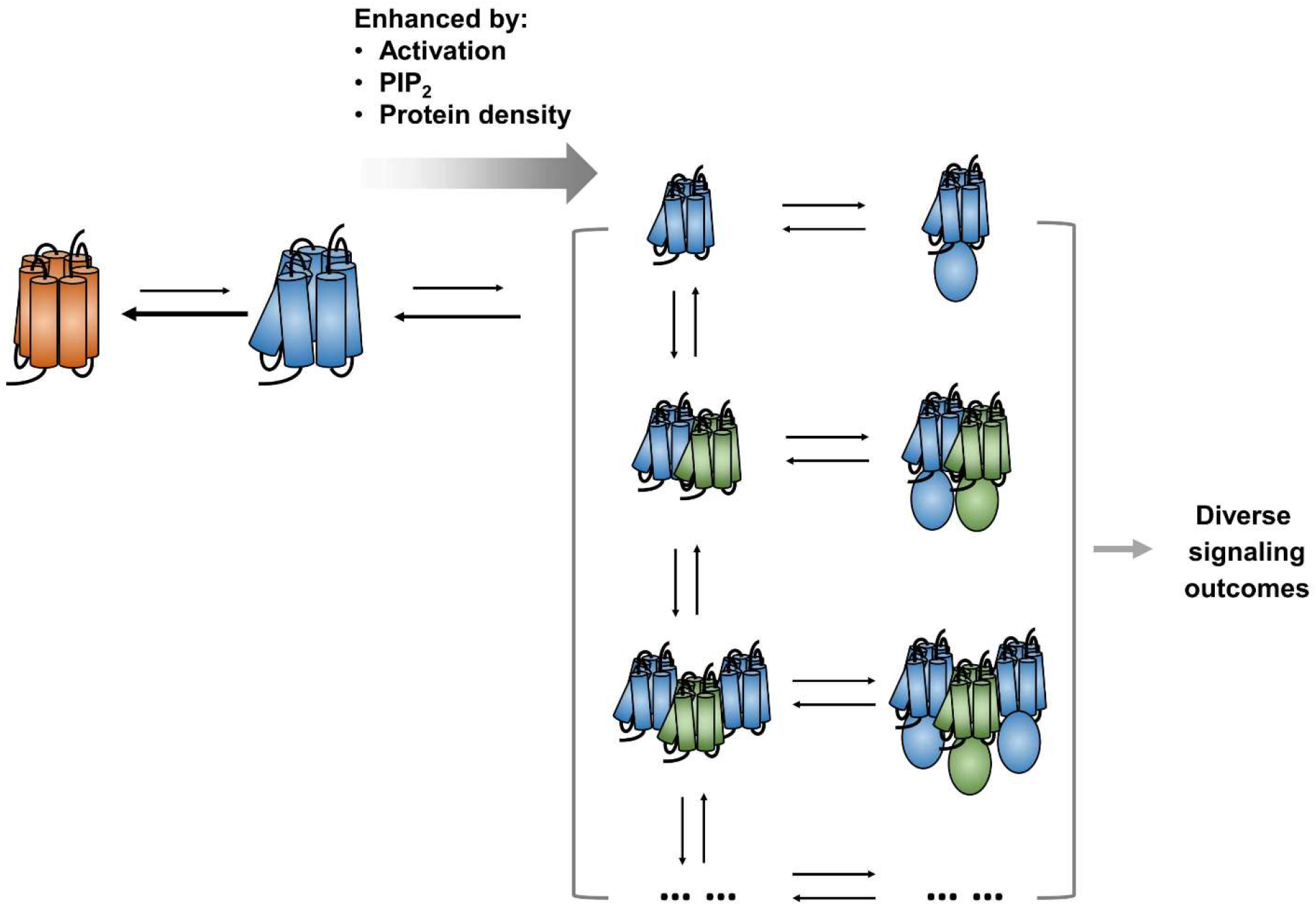
Allosteric modulation and GPCR oligomerisation. The activated GPCR generates an array of oligomeric assemblies which couples to signalling partners, generating an array of supramolecular signalling complexes capable of initiating different signalling pathways. This mechanism of combinatory modulation of GPCR is responsive to the lipid bilayer environment and the conformational state of the receptor. Receptor activation, PIP_2_ and/or mini-Gs binding, and an increase in receptor density in the bilayer all promote oligomerisation.

## Methods

### MD simulation set-up

Simulations used the MARTINI coarse-grained forcefield (*67*) as previously applied in simulations of monomeric A2aR (*37*). Nine or sixteen copies of the receptor in a given conformational state were individually rotated through a random angle at the centre of a box of size of 45 × 45 × 25 nm^3^ and then randomly translated in the *xy* plane. Both randomization processes were carried out by the random module of NumPy. An even distribution was made sure in the starting configurations. The resultant configuration was embedded in a membrane bilayer with the specified complex lipid composition (see Table 1) using the *insane.py* script(*68*). Electrolyte solution corresponding to ~0.15 M NaCl was added and additional sodium ions were added to neutralise the system. All the simulations were performed using GROMACS 5.1(*69*). The CG simulations parameters were taken from ref (*37*). A summary of simulations performed is provided in Table 1, amounting to a total of ~2.6 ms of simulation data collected.

### Characterisation of oligomeric configurations

The minimum distance between each pair of proteins was monitored as a function of time (SI Fig. S1), from which the density of such distance distribution was estimated. The density estimation showed a first peak at ~0.55 nm and a first trough at ~0.7 nm. We found that a cut-off of 0.75 nm could best discriminate effective associations. Using this cut-off and hierarchical clustering, the number of oligomers of different orders were counted and each copy of the receptor was assigned to those identified oligomers separately for each frame. The identified oligomeric assemblies with the same oligomeric order were grouped together into an ‘oligomer pool’, and characterised, *i.e.* clustered, within each pool if the order is lower than 6. Since calculation of root mean square deviation (RMSD) based on structural coordinates is sensitive to the order of the receptor indexing, we set out to assign an order to the receptor copies in the oligomers. We took a reference structure randomly from each oligomer pool and calculated the coordinate RMSDs for the rest of the structures in that pool, testing all possible orders of the receptor copies. The order giving the lowest RMSD was assigned to the structure. The oligomeric structures in each oligomer pool were then clustered using the KMeans method based on their reordered coordinates. The number of clusters were manually tested for each oligomer pool to ensure a homogeneous distribution of structures in each cluster. These clusters were individually labelled as *A.b* where *A* was the oligomeric order and *b* the cluster identifier in their oligomer pool. To assist structural inspection, the coarse-grained oligomer models were converted back to atomistic models using CHARMM 36 force field and the *backward.py* script provided by MARTINI website. A repository of the 9-copy oligomer atomistic models can be found at http://doi.org/10.5281/zenodo.4300676.

### Calculation of oligomer residence time

For every oligomer cluster, the durations of their continuous appearances in the simulations were recorded. Of note, visual inspection of trajectories revealed that protein-protein interactions can have quick flickering in contact distance that does not lead to ‘real’ dissociations in MARTINI coarse-grained force field. Hence, we treated the protein-protein associations as ‘being continuous’ if their inter-protomer contacts broke for a time period shorter than 100 ns. The oligomer residence time was calculated from the normalised survival time-correlation function *σ*(*t*):

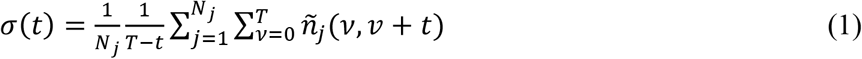

where *T* is the total simulation time, N_j_ the number of continuous appearances collected from simulations, and 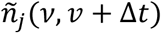 is a function that takes the value 1 if the oligomer appeared for a continuous duration of Δ*t* after forming the contact at time *v*, and takes the value 0 if otherwise. Bi exponentials 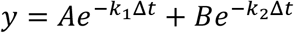 were used to fit the survival curve (Fig 3A). The smaller *k* was regarded as oligomer *k*_*off*_ and the oligomer residence time was calculated as 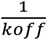.

### Characterisation of lipid interactions

The lipid binding sites, and binding kinetics were calculated using PyLipID (https://github.com/wlsong/PyLipID). For lipid interactions with a given residue, a continuous lipid contact was defined using a dual cut-off scheme such that the interaction started when lipid headgroup moved to a given residue closer than 0.55 nm and ended when the headgroup moved farther than 1.0 nm. The residue-wise lipid residence time was calculated from the durations of these continuous contacts using function (*1*) where *N*_*j*_ the number of continuous lipid contacts with the residue collected from simulations. Lipid binding sites were derived from the lipid contact information using graph theory and community analysis. Graphs were constructed using residues as nodes and the frequency with which a given pair of residues interacted with the same lipid molecule as edges. The best partition function of the community library (https://github.com/taynaud/python-louvain) was used to detect community structures. Each calculated community was considered a binding site. A continuous lipid contact with a given binding site starts as the lipid headgroup moved closer than 0.55 nm to any of the residues in the binding site and ends as the headgroup moved farther than 1.0 nm. Similar to the residue-wise residence time, the lipid residence time of a given binding site was calculated from the durations of these continuous binding site contacts using function (*1*) where *N*_*j*_ the number of continuous lipid contacts with the binding site collected from simulations. The analysis data of PIP_2_ and DOPS interactions with A2aR can be found at https://github.com/wlsong/LipidInteractions.

### MSM and validation

Markov state models were used to understand A2aR oligomerisation network. Since the oligomerisation process of a receptor is dependent on the state of the rest of the membrane, we built MSMs using the oligomerisation state of the membrane. The number of oligomers of various orders were counted at every simulation frame to represent the oligomerisation state of the membrane, which were labelled as *A*^*a*^-*B*^*b*^-*C*^*c*^ in which *A, B, C* denote the oligomeric orders and *a, b, c* represent the number of oligomers of the denoting order in the membrane. The transitions between these oligomerisation states were counted and the transition matrix *P*_*ij*_ which mapped out the probability of transitioning from state *I* at time *t* to state *j* at time *t* + *τ* were estimated using Maximum likelihood estimation (*70*). The transition matrix was calculated using pyEMMA (*42*). The Markovian behaviour of the process can be checked by plotting the implied timescales (*k*) from eigenvalues *μ* of the transition matrix *P* at various lag times *τ* based on the relationship *k* = *− τ/ln(μ)*. The smallest *τ*, at which the implied timescales started to converge to an unchanged rate, was taken to build the MSM. *τ* = 5 *μS* was used for both 9-copy systems and 9-copy NoPIP_2_ systems. The mean first passage time (mfpt) between the macrostates was calculated as described in ref (*71*) via the functionality in PyEMMA. The lifetime of an oligomerisation state S_i_ was calculated as mfpt(*i,j*), where *j* ∈ Θ and *j* ≠ *i*.

## Acknowledgements

Research in the M.S.P.S. group is supported by Wellcome (208361/Z/17/Z), BBSRC (BB/R00126X/1), and PRACE (Partnership for Advanced Computing in Europe; 2016163984). W.S. acknowledges support from the Newton International Fellowship. This project made use of time on ARCHER via the HECBioSim, supported by EPSRC (EP/L000253/1).

## References

1. A. Glukhova, C. J. Draper-Joyce, R. K. Sunahara, A. Christopoulos, D. Wootten, P. M. Sexton, Rules of engagement: GPCRs and G proteins, ACS Pharmacol. Transl. Sci. 1, 73–83 (2018).

2. G. Milligan, R. J. Ward, S. Marsango, GPCR homo-oligomerization, Curr. Opin. Cell Biol. 57, 40–47 (2019).

3. W. Guo, E. Urizar, M. Kralikova, J. C. Mobarec, L. Shi, M. Filizola, J. A. Javitch, Dopamine D2 receptors form higher order oligomers at physiological expression levels, EMBO J. 27, 2293–2304 (2008).

4. L. F. Pisterzi, D. B. Jansma, J. Georgiou, M. J. Woodside, J. T. C. Chou, S. Angers, V. Raicu, J. W. Wells, Oligomeric Size of the M2 muscarinic receptor in live cells as determined by quantitative fluorescence resonance energy transfer, J. Biol. Chem. 285, 16723–16738 (2010).

5. S. M. Walsh, S. Mathiasen, S. M. Christensen, J. F. Fay, C. King, D. Provasi, E. Borrero, S. G. F. Rasmussen, J. J. Fung, M. Filizola, K. Hristova, B. Kobilka, D. L. Farrens, D. Stamou, Single proteoliposome high-content analysis reveals differences in the homo-oligomerization of GPCRs, Biophys. J. 115, 300–312 (2018).

6. R. J. Ward, J. D. Pediani, A. G. Godin, G. Milligan, Regulation of oligomeric organization of the serotonin 5-hydroxytryptamine 2C (5-HT2C) receptor observed by spatial intensity distribution analysis, J Biol Chem 290, 12844–12857 (2015).

7. A. Tabor, S. Weisenburger, A. Banerjee, N. Purkayastha, J. M. Kaindl, H. Hübner, L. Wei, T. W. Grömer, J. Kornhuber, N. Tschammer, N. J. M. Birdsall, G. I. Mashanov, V. Sandoghdar, P. Gmeiner, Visualization and ligand-induced modulation of dopamine receptor dimerization at the single molecule level, Sci. Reports 6, 33233 (2016).

8. R. J. Ward, J. D. Pediani, K. G. Harikumar, L. J. Miller, G. Milligan, Spatial intensity distribution analysis quantifies the extent and regulation of homodimerization of the secretin receptor, Biochem J 474, 1879–1895 (2017).

9. J. Paprocki, G. Biener, M. Stoneman, V. Raicu, In-cell detection of conformational substates of a G protein-coupled receptor quaternary structure: modulation of substate probability by cognate ligand binding, J. Phys. Chem. B 10.1021/acs.jpcb.0c06081, DOI 10.1021/acs.jpcb.1020c06081 (2020).

10. M. R. Stoneman, G. Biener, R. J. Ward, J. D. Pediani, D. Badu, A. Eis, I. Popa, G. Milligan, V. Raicu, A general method to quantify ligand-driven oligomerization from fluorescence-based images, Nature Methods 16, 493–496 (2019).

11. R. V. Shivnaraine, B. Kelly, K. S. Sankar, D. S. Redka, Y. R. Han, F. Huang, G. Elmslie, D. Pinto, Li, J. V. Rocheleau, C. C. Gradinaru, J. Ellis, J. W. Wells, Allosteric modulation in monomers and oligomers of a G protein-coupled receptor, Elife 5, e11685 (2016).

12. T. Buenaventura, S. Bitsi, W. E. Laughlin, T. Burgoyne, Z. Lyu, A. I. Oqua, H. Norman, E. R. McGlone, A. S. Klymchenko, I. R. Corrêa, Jr., A. Walker, A. Inoue, A. Hanyaloglu, J. Grimes, Koszegi, D. Calebiro, G. A. Rutter, S. R. Bloom, B. Jones, A. Tomas, Agonist-induced membrane nanodomain clustering drives GLP-1 receptor responses in pancreatic beta cells, PLOS Biology 17, e3000097 (2019).

13. B. Ge, J. Lao, J. Li, Y. Chen, Y. Song, F. Huang, Single-molecule imaging reveals dimerization/oligomerization of CXCR4 on plasma membrane closely related to its function, Sci Rep 7, 16873 (2017).

14. J. Möller, A. Isbilir, T. Sungkaworn, B. Osberg, C. Karathanasis, V. Sunkara, E. O. Grushevskyi, A. Bock, P. Annibale, M. Heilemann, C. Schütte, M. J. Lohse, Single-molecule analysis reveals agonist-specific dimer formation of μ-opioid receptors, Nature Chem. Biol. 16, 946–954 (2020).

15. G. Navarro, A. Cordomí, M. Zelman-Femiak, M. Brugarolas, E. Moreno, D. Aguinaga, L. Perez-Benito, A. Cortés, V. Casadó, J. Mallol, E. I. Canela, C. Lluís, L. Pardo, A. J. García-Sáez, P. J. McCormick, R. Franco, Quaternary structure of a G-protein-coupled receptor heterotetramer in complex with Gi and Gs, BMC Biology 14, 26 (2016).

16. R. V. Shivnaraine, D. D. Fernandes, H. Ji, Y. Li, B. Kelly, Z. Zhang, Y. R. Han, F. Huang, K. S. Sankar, D. N. Dubins, J. V. Rocheleau, J. W. Wells, C. C. Gradinaru, Single-molecule analysis of the supramolecular organization of the M2 muscarinic receptor and the Gαi1 protein, J. Amer. Chem. Soc. 138, 11583–11598 (2016).

17. H. Yang, L. Qu, J. Ni, M. Wang, Y. Huang, Palmitoylation participates in G protein coupled signal transduction by affecting its oligomerization, Molec. Membrane Biol. 25, 58–71 (2008).

18. Y. Li, R. V. Shivnaraine, F. Huang, J. W. Wells, C. C. Gradinaru, Ligand-induced coupling between oligomers of the M2 Receptor and the Gi1 protein in live cells, Biophys. J. 115, 881–895 (2018).

19. V. Cherezov, D. M. Rosenbaum, M. A. Hanson, S. G. F. Rasmussen, F. S. Thian, T. S. Kobilka, H.-J. Choi, P. Kuhn, W. I. Weis, B. K. Kobilka, R. C. Stevens, High-resolution crystal structure of an engineered human β2-adrenergic G protein–coupled receptor, Science 318, 1258–1265 (2007).

20. A. Manglik, A. C. Kruse, T. S. Kobilka, F. S. Thian, J. M. Mathiesen, R. K. Sunahara, L. Pardo, W. I. Weis, B. K. Kobilka, S. Granier, Crystal structure of the micro-opioid receptor bound to a morphinan antagonist, Nature 485, 321–326 (2012).

21. W. Huang, A. Manglik, A. J. Venkatakrishnan, T. Laeremans, E. N. Feinberg, A. L. Sanborn, H. E. Kato, K. E. Livingston, T. S. Thorsen, R. C. Kling, S. Granier, P. Gmeiner, S. M. Husbands, J. R. Traynor, W. I. Weis, J. Steyaert, R. O. Dror, B. K. Kobilka, Structural insights into μ-opioid receptor activation, Nature 524, 315–321 (2015).

22. B. Wu, E. Y. Chien, C. D. Mol, G. Fenalti, W. Liu, V. Katritch, R. Abagyan, A. Brooun, P. Wells, F. C. Bi, D. J. Hamel, P. Kuhn, T. M. Handel, V. Cherezov, R. C. Stevens, Structures of the CXCR4 chemokine GPCR with small-molecule and cyclic peptide antagonists, Science 330, 1066–1071 (2010).

23. K. Zhang, J. Zhang, Z.-G. Gao, D. Zhang, L. Zhu, G. W. Han, S. M. Moss, S. Paoletta, E. Kiselev, W. Lu, G. Fenalti, W. Zhang, C. E. Müller, H. Yang, H. Jiang, V. Cherezov, V. Katritch, K. A. Jacobson, R. C. Stevens, B. Wu, Q. Zhao, Structure of the human P2Y12 receptor in complex with an antithrombotic drug, Nature 509, 115 (2014).

24. W. Liu, E. Chun, A. A. Thompson, P. Chubukov, F. Xu, V. Katritch, G. W. Han, C. B. Roth, L. H. Heitman, A. P. IJzerman, V. Cherezov, R. C. Stevens, Structural basis for allosteric regulation of GPCRs by sodium ions, Science 337, 232–236 (2012).

25. D. Zhang, Z. G. Gao, K. Zhang, E. Kiselev, S. Crane, J. Wang, S. Paoletta, C. Yi, L. Ma, W. Zhang, G. W. Han, H. Liu, V. Cherezov, V. Katritch, H. Jiang, R. C. Stevens, K. A. Jacobson, Q. Zhao, B. Wu, Two disparate ligand-binding sites in the human P2Y1 receptor, Nature 520, 317–321 (2015).

26. A. L. Duncan, W. Song, M. S. P. Sansom, Lipid-dependent regulation of ion channels and GPCRs: insights from structures and simulations, Ann. Rev. Pharmacol. Toxicol. 60, 31–50 (2020).

27. S. Gahbauer, R. A. Böckmann, Membrane-mediated oligomerization of G protein coupled receptors and its implications for GPCR function, Frontiers Physiol. 7, 494 (2016).

28. X. Periole, A. M. Knepp, T. P. Sakmar, S. J. Marrink, T. Huber, Structural determinants of the supramolecular organization of G protein-coupled receptors in bilayers, J. Amer. Chem. Soc. 134, 10959–10965 (2012).

29. K. Pluhackova, S. Gahbauer, F. Kranz, T. A. Wassenaar, R. A. Bockmann, Dynamic cholesterol-conditioned dimerization of the G protein coupled chemokine receptor type 4, PLoS Comp. Biol. 12, e1005169 (2016).

30. X. Prasanna, D. Sengupta, A. Chattopadhyay, Cholesterol-dependent conformational plasticity in GPCR dimers, Sci. Reports 6, 31858 (2016).

31. D. Provasi, M. B. Boz, J. M. Johnston, M. Filizola, Preferred supramolecular organization and dimer interfaces of opioid receptors from simulated self-association, PLoS Comp. Biol. 11, e1004148 (2015).

32. D. Meral, D. Provasi, D. Prada-Gracia, J. Moller, K. Marino, M. J. Lohse, M. Filizola, Molecular details of dimerization kinetics reveal negligible populations of transient mu-opioid receptor homodimers at physiological concentrations, Sci. Reports 8, 7705 (2018).

33. P. A. Vidi, B. R. Chemel, C. D. Hu, V. J. Watts, Ligand-dependent oligomerization of dopamine D2 and adenosine A(2A) receptors in living neuronal cells, Molec. Pharmacol. 74, 544–551 (2008).

34. P. A. Vidi, J. J. Chen, J. M. K. Irudayaraj, V. J. Watts, Adenosine A(2A) receptors assemble into higher-order oligomers at the plasma membrane, FEBS Lett. 582, 3985–3990 (2008).

35. S. J. Marrink, J. Risselada, S. Yefimov, D. P. Tieleman, A. H. de Vries, The MARTINI force field: coarse grained model for biomolecular simulations, J. Phys. Chem. B. 111, 7812–7824 (2007).

36. L. Monticelli, S. K. Kandasamy, X. Periole, R. G. Larson, D. P. Tieleman, S. J. Marrink, The MARTINI coarse grained force field: extension to proteins, J. Chem. Theor. Comp. 4, 819–834 (2008).

37. W. L. Song, H. Y. Yen, C. V. Robinson, M. S. P. Sansom, State-dependent lipid interactions with the A2a receptor revealed by MD simulations using *in vivo*-mimetic membranes, Structure 27, 392–403 (2019).

38. H. Y. Yen, K. K. Hoi, I. Liko, G. Hedger, M. R. Horrell, W. L. Song, D. Wu, P. Heine, T. Warne, Y. Lee, B. Carpenter, A. Pluckthun, C. G. Tate, M. S. P. Sansom, C. V. Robinson, PtdIns(4,5)P-2 stabilizes active states of GPCRs and enhances selectivity of G-protein coupling, Nature 559, 424–427 (2018).

39. W. Huang, M. Masureel, Q. Qianhui, J. Janetzko, A. Inoue, H. E. Kato, M. J. Robertson, K. C. Nguyen, J. S. Glenn, G. Skiniotis, B. K. Kobilka, Structure of the neurotensin receptor 1 in complex with β-arrestin 1, Nature 579, 303–308 (2020).

40. S. Gahbauer, K. Pluhackova, R. A. Bockmann, Closely related, yet unique: Distinct homo- and heterodimerization patterns of G protein coupled chemokine receptors and their fine-tuning by cholesterol, PLoS Comput Biol 14, e1006062 (2018).

41. B. E. Husic, V. S. Pande, Markov state models: from an art to a science, J. Amer. Chem. Soc. 140, 2386–2396 (2018).

42. M. K. Scherer, B. Trendelkamp-Schroer, F. Paul, G. Perez-Hernandez, M. Hoffmann, N. Plattner, C. Wehmeyer, J. H. Prinz, F. Noe, PyEMMA 2: A software package for estimation, validation, and analysis of Markov models, J. Chem. Theor. Comput. 11, 5525–5542 (2015).

43. G. Navarro, S. Ferre, A. Cordomi, E. Moreno, J. Mallol, V. Casado, A. Cortes, H. Hoffmann, J. Ortiz, E. I. Canela, C. Lluis, L. Pardo, R. Franco, A. S. Woods, Interactions between intracellular domains as key determinants of the quaternary structure and function of receptor heteromers, J. Biol. Chem. 285, 27346–27359 (2010).

44. V. Velazhahan, N. Ma, G. Pándy-Szekeres, A. J. Kooistra, Y. Lee, D. E. Gloriam, N. Vaidehi, C. G. Tate, Structure of the class D GPCR Ste2 dimer coupled to two G proteins, Nature 10.1038/s41586-020-2994-1 (2020).

45. J. Wang, L. He, C. A. Combs, G. Roderiquez, M. A. Norcross, Dimerization of CXCR4 in living malignant cells: control of cell migration by a synthetic peptide that reduces homologous CXCR4 interactions, Molec. Cancer Therapeut. 5, 2474–2483 (2006).

46. R. Guixà-González, M. Javanainen, M. Gómez-Soler, B. Cordobilla, J. C. Domingo, F. Sanz, M. Pastor, F. Ciruela, H. Martinez-Seara, J. Selent, Membrane omega-3 fatty acids modulate the oligomerisation kinetics of adenosine A2A and dopamine D2 receptors, Scientific Reports 6, 19839 (2016).

47. D. Calebiro, T. Sungkaworn, Single-molecule imaging of GPCR interactions, Trends Pharmacol. Sci. 39, 109–122 (2018).

48. G. van den Bogaart, K. Meyenberg, H. J. Risselada, H. Amin, K. I. Willig, B. E. Hubrich, M. Dier, S. W. Hell, H. Grubmüller, U. Diederichsen, R. Jahn, Membrane protein sequestering by ionic protein-lipid interactions, Nature 479, 552–555 (2011).

49. K. D. Nguyen, M. Vigers, E. Sefah, S. Seppala, J. P. Hoover, N. S. Schonenbach, B. Mertz, M. A. O’Malley, S. Han, Oligomerization of the human adenosine A2A receptor is driven by the intrinsically disordered C-Terminus, BioRxiv 10.1101/2020.12.21.423144 %J bioRxiv, 2020.2012.2021.423144 (2020).

50. L. Tovo-Rodrigues, A. Roux, M. H. Hutz, L. A. Rohde, A. S. Woods, Functional characterization of G-protein-coupled receptors: a bioinformatics approach, Neuroscience 277, 764–779 (2014).

51. R. S. Kasai, T. K. Fujiwara, A. Kusumi, Metastable GPCR dimers trigger the basal signal by recruiting G-proteins, BioRxiv 10.1101/2020.02.10.929588 %J bioRxiv, 2020.2002.2010.929588 (2020).

52. M. Patrone, E. Cammarota, V. Berno, P. Tornaghi, D. Mazza, M. Degano, Combinatorial allosteric modulation of agonist response in a self-interacting G-protein coupled receptor, Comms. Biol. 3, 27 (2020).

53. X. Periole, T. Huber, S. J. Marrink, T. P. Sakmar, G protein-coupled receptors self-assemble in dynamics simulations of model bilayers, J. Amer. Chem. Soc. 129, 10126–10132 (2007).

54. M. Chavent, A. P. Chetwynd, P. J. Stansfeld, M. S. Sansom, Dimerization of the EphA1 receptor tyrosine kinase transmembrane domain: Insights into the mechanism of receptor activation, Biochem. 53, 6641–6652 (2014).

55. J. Domanski, M. S. P. Sansom, P. J. Stansfeld, R. B. Best, Balancing force field protein-lipid interactions to capture transmembrane helix-helix association, J. Chem. Theor. Comput. 14, 1706–1715 (2018).

56. S. Gahbauer, R. A. Bockmann, Comprehensive characterization of lipid-guided G protein-coupled receptor dimerization, J. Phys. Chem. B 124, 2823–2834 (2020).

57. X. Periole, T. Zeppelin, B. Schiott, Dimer interface of the human serotonin transporter and effect of the membrane composition, Sci. Reports 8, 5080 (2018).

58. G. Hedger, M. S. P. Sansom, Lipid interaction sites on channels, transporters and receptors: recent insights from molecular dynamics simulations, Biochim. Biophys. Acta 1858, 2390–2400 (2016).

59. V. Corradi, E. Mendez-Villuendas, H. I. Ingólfsson, R.-X. Gu, I. Siuda, M. N. Melo, A. Moussatova, L. J. DeGagné, B. I. Sejdiu, G. Singh, T. A. Wassenaar, K. Delgado Magnero, S. J. Marrink, D. P. Tieleman, Lipid–protein interactions are unique fingerprints for membrane proteins, ACS Central Sci. 4, 709–717 (2018).

60. A. L. Duncan, R. A. Corey, M. S. P. Sansom, Defining how multiple lipid species interact with inward rectifier potassium (Kir2) channels, Proc. Natl. Acad. Sci. USA 117, 7803–7813 (2020).

61. M. Javanainen, H. Martinez-Seara, I. Vattulainen, Excessive aggregation of membrane proteins in the Martini model, Plos One 12, e0187936 (2017).

62. R. Alessandri, P. C. Telles de Souza, S. Thallmair, M. N. Melo, A. H. De Vries, S. J. Marrink, Pitfalls of the Martini model, J. Chem. Theor. Comput. 15, 5448–5460 (2019).

63. G. Navarro, A. Cordomí, V. Casadó-Anguera, E. Moreno, N.-S. Cai, A. Cortés, E. I. Canela, C. W. Dessauer, V. Casadó, L. Pardo, C. Lluís, S. Ferré, Evidence for functional pre-coupled complexes of receptor heteromers and adenylyl cyclase, Nature Comms. 9, 1242 (2018).

64. P. Hein, M. Frank, C. Hoffmann, M. J. Lohse, M. Bünemann, Dynamics of receptor/G protein coupling in living cells, EMBO J. 24, 4106–4114 (2005).

65. B. H. Falkenburger, J. B. Jensen, B. Hille, Kinetics of M1 muscarinic receptor and G protein signaling to phospholipase C in living cells, J. Gen. Physiol. 135, 81–97 (2010).

66. S. Ferre, V. Casado, L. A. Devi, M. Filizola, R. Jockers, M. J. Lohse, G. Milligan, J. P. Pin, X. Guitart, G protein-coupled receptor oligomerization revisited: Functional and pharmacological perspectives, Pharmacol. Rev. 66, 413–434 (2014).

67. D. H. de Jong, G. Singh, W. F. D. Bennett, C. Arnarez, T. A. Wassenaar, L. V. Schäfer, X. Periole, D. P. Tieleman, S. J. Marrink, Improved parameters for the Martini coarse-grained protein force field, J. Chem. Theor. Comput. 9, 687–697 (2013).

68. T. A. Wassenaar, H. I. Ingólfsson, R. A. Böckmann, D. P. Tieleman, S. J. Marrink, Computational lipidomics with insane: a versatile tool for generating custom membranes for molecular simulations, J. Chem. Theor. Comput. 11, 2144–2155 (2015).

69. M. J. Abraham, T. Murtola, R. Schulz, S. Páll, J. C. Smith, B. Hess, E. Lindahl, GROMACS: High performance molecular simulations through multi-level parallelism from laptops to supercomputers, SoftwareX 1-2, 19–25 (2015).

70. J.-H. Prinz, H. Wu, M. Sarich, B. Keller, M. Senne, M. Held, J. D. Chodera, C. Schütte, F. Noé, Markov models of molecular kinetics: Generation and validation, J. Chem. Phys. 134, 174105 (2011).

71. N. Singhal, V. S. Pande, Error analysis and efficient sampling in Markovian state models for molecular dynamics, J. Chem. Phys. 123, 204909 (2005).

