## Supplementary material for "Modulation of A2aR Oligomerisation by Conformational State and PIP_2_ Interactions Revealed by MD Simulations and Markov Models": SI Figures & Tables

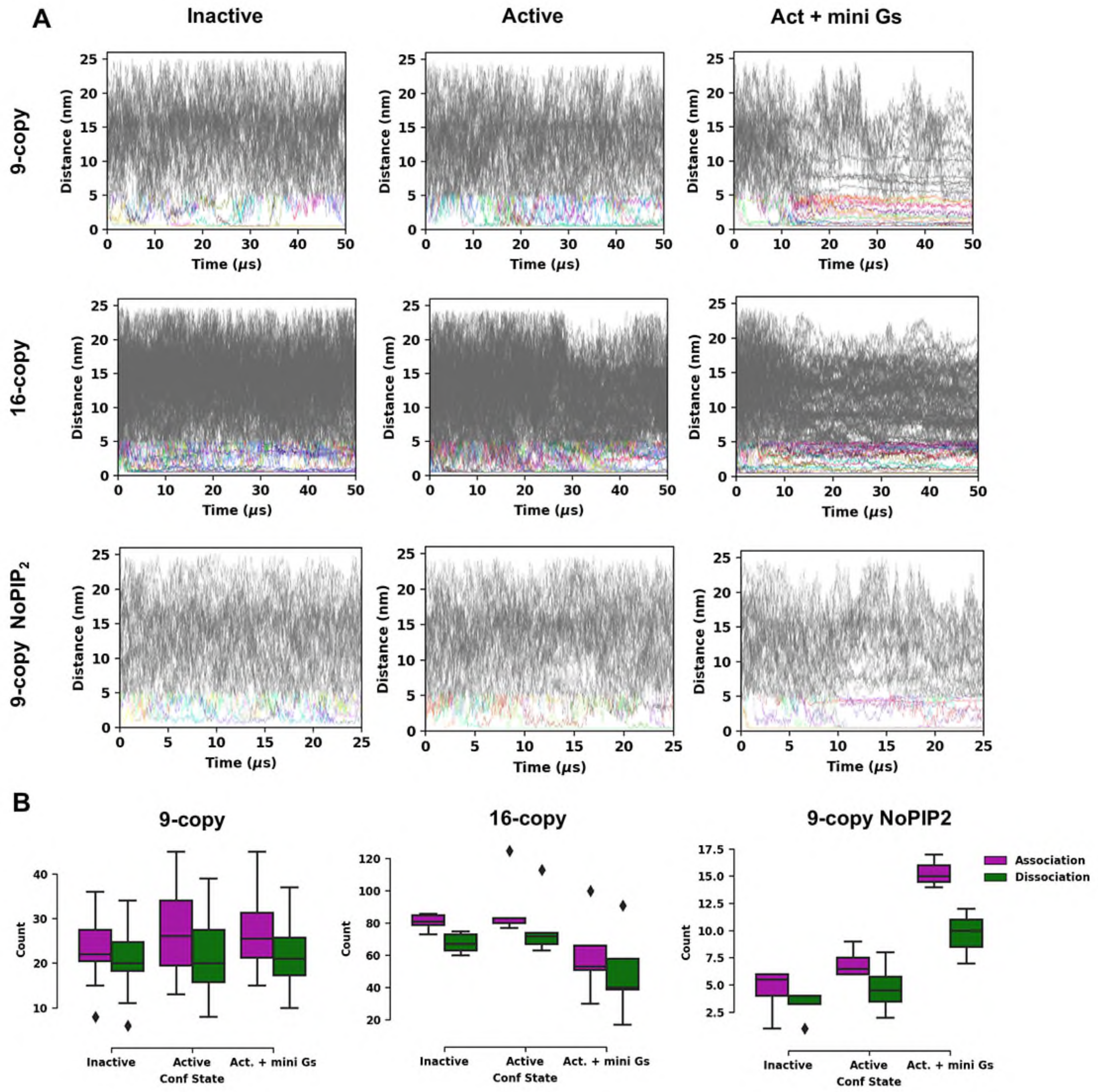


**SI Figure S1. Associations and dissociations sampled in simulations.** (A) Pair-wise minimum distance vs. time is shown for one example of a trajectory from each simulation ensemble. The minimum distance between each pair of proteins in the selected trajectory is plotted as a function of time. Distances below 5 nm are differentiated using different colours and coloured in grey when above 5 nm. (B) The number of associations (defined as a pair of proteins coming closer than 0.75 nm) and dissociations (defined as a pair of proteins separating to further than 0.75 nm) sampled in each trajectory of the simulation ensembles. Standard box plot settings were used, *i.e.* the boxes covering the interquartile range (IQR) between the lower and upper quartiles and whiskers showing a distance of 1.5 IQR above the upper quartile.

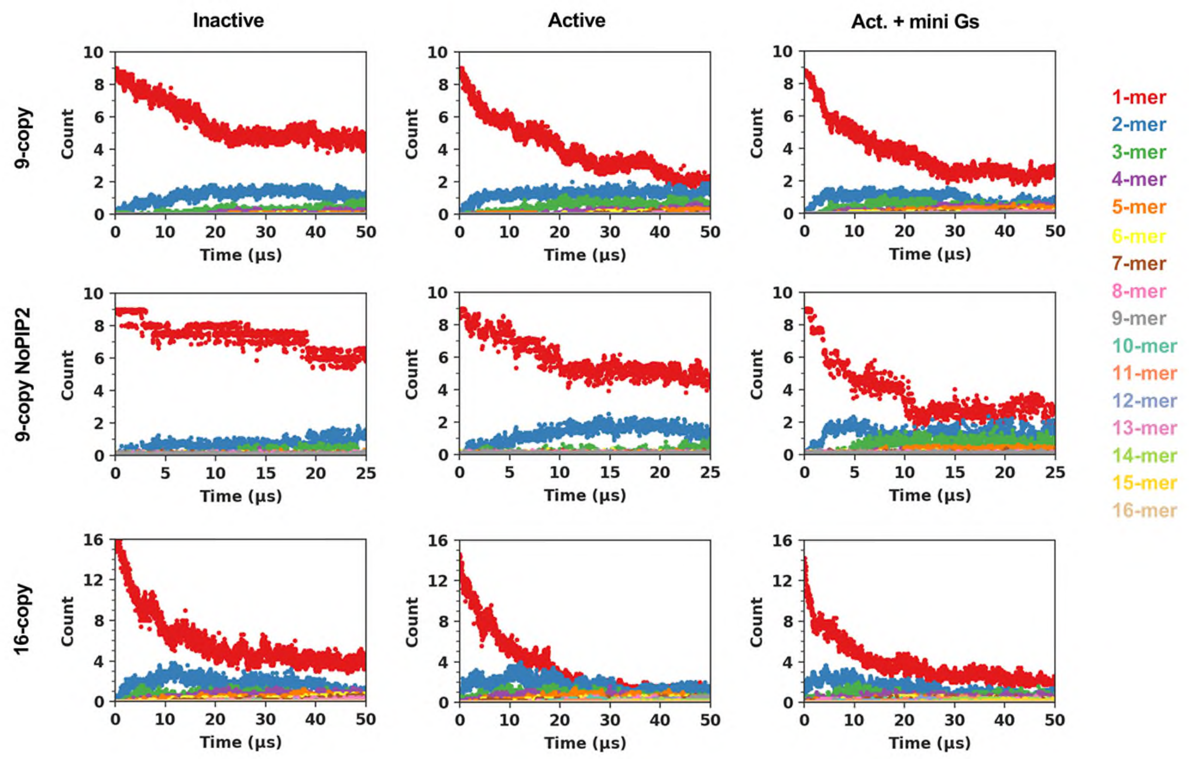


**SI Figure S2. Time course of oligomer formation.** All Data points across trajectories in each simulation ensemble were plotted together in each panel.

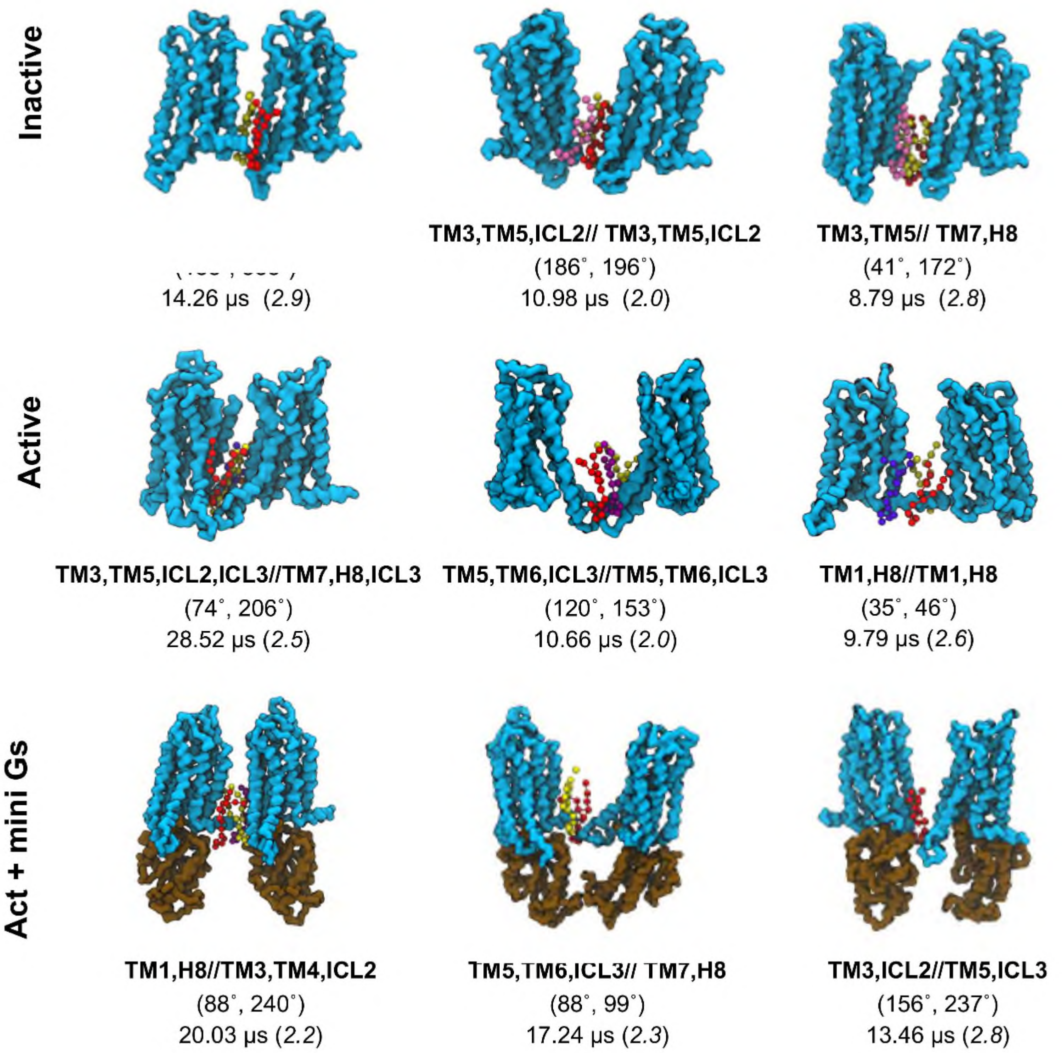


**SI Figure S3. Stable dimeric configurations from 9-copy simulations.** The 3 dimeric configurations with the longest residence time were shown for each conformational state. The backbone beads of the receptor are shown in cyan surface and those of mini Gs in brown surface. The $PIP_2$ molecules at the association interfaces are shown in ball and sticks with different colours. Below each of the structures are the association interfaces in bold, the average values of the binding angles, and the dimer residence time which is followed by the cluster id label in brackets. A table of dimeric configuration profiles can be found in SI Table S1. The atomistic models of the dimeric configurations can be downloaded at http://doi.org/10.5281/zenodo.4300676.

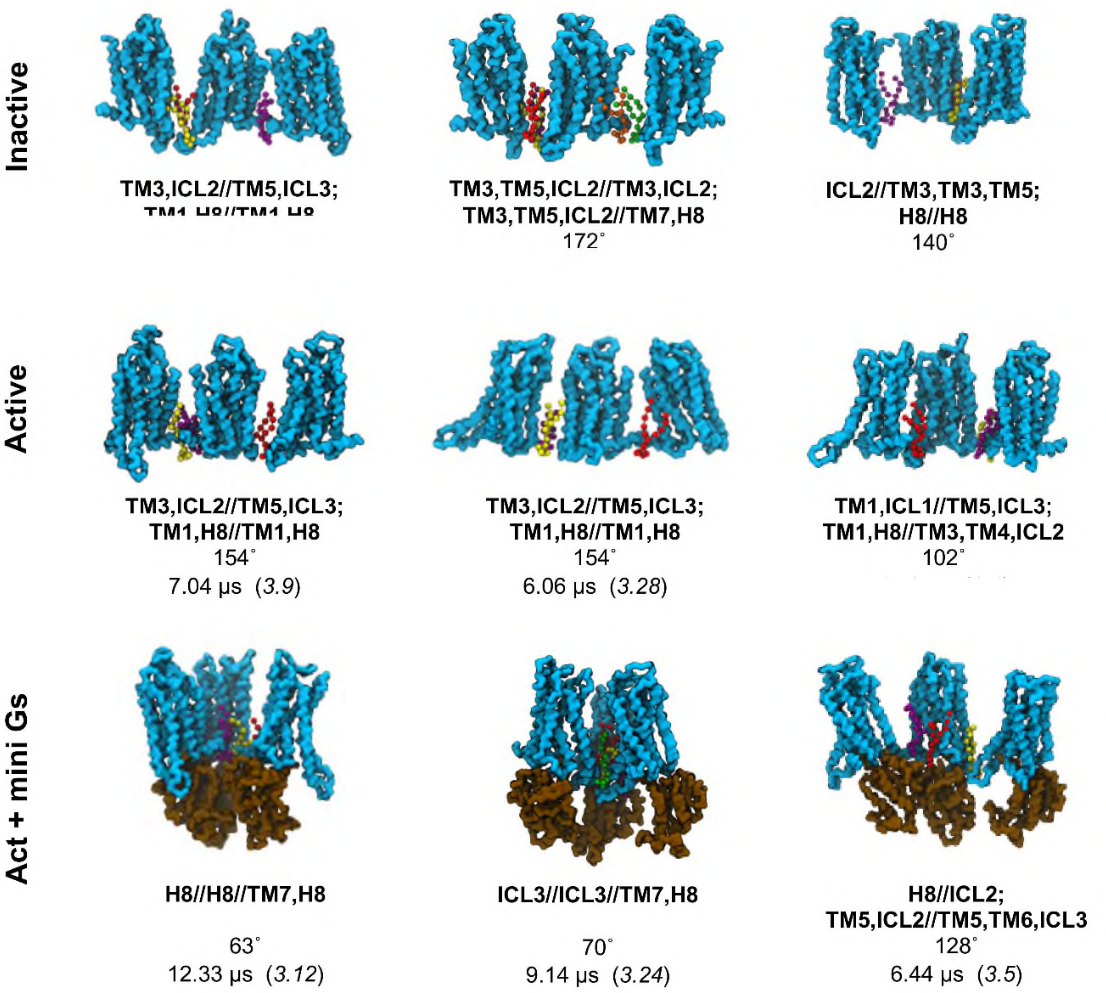


**SI Figure S4. Stable trimer configurations from 9-copy simulations.** The 3 trimeric configurations with the longest residence time were shown for each conformational state. The backbone beads of the receptor are shown in cyan surface and those of mini Gs in brown surface. The $PIP_2$ molecules at the association interfaces are shown in ball and sticks with different colours. Below each of the structures are the association interfaces in bold, the average values of the binding angles, and the trimer residence time which is followed by the cluster id label in brackets. A table of trimeric configuration profiles together with the atomistic models of the trimeric configurations can be downloaded at http://doi.org/10.5281/zenodo.4300676.

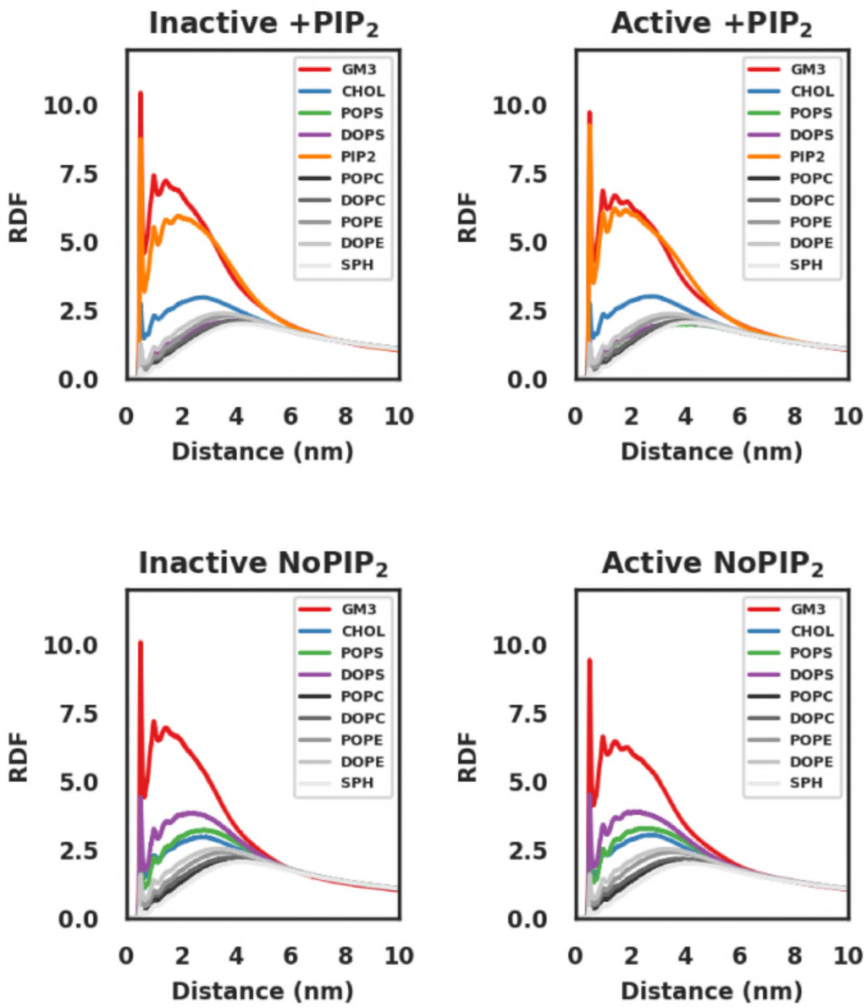


**SI Figure S5. Lipid Radial distribution function around the receptor** in the 9-copy simulations (+$PIP_2$) and the 9-copy NoPIP$_2$ simulations.

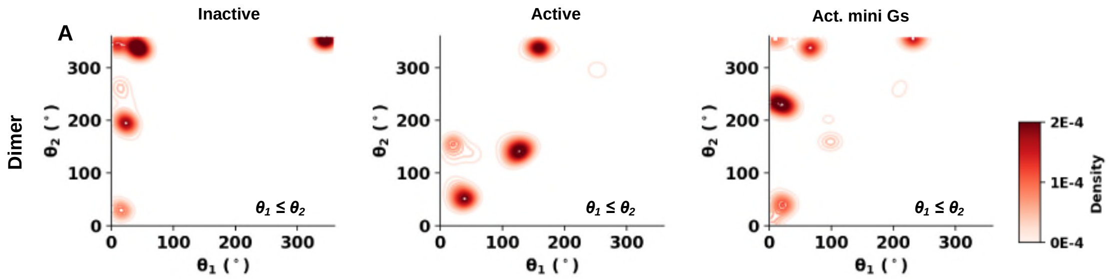


**SI Figure S6. Dimer binding angles in 9-copy NoPIP2 simulations.** The binding angle is defined as the angle between dimer binding interface and the principal axis that is parallel to H8, as illustrated in Fig 3B.

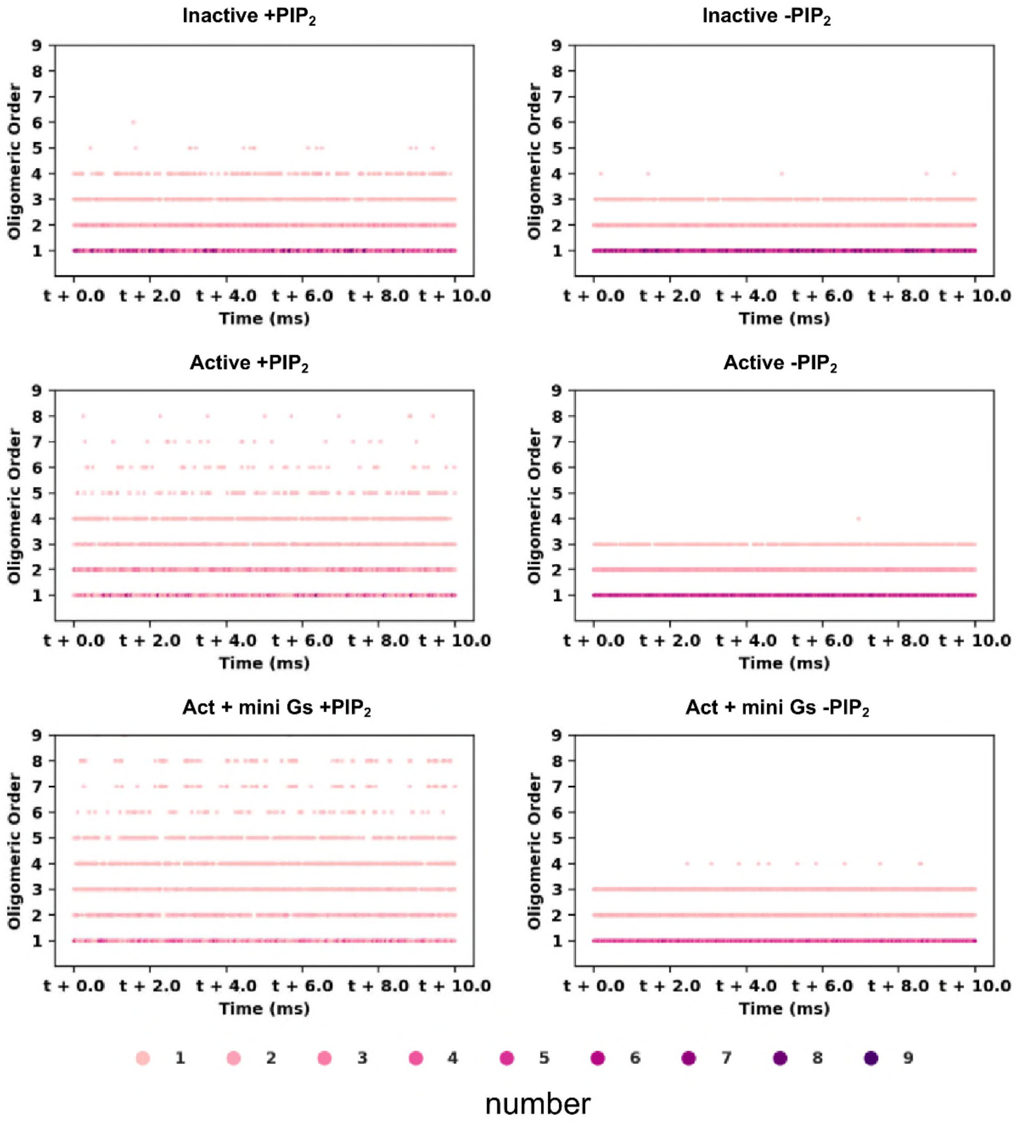


**SI Figure S7. Oligomerisation kinetics in the 9-copy systems.** A 10 ms trajectory was generated for each of the conformational state in 9-copy (+$PIP_2$) and 9-copy NoPIP2 (-$PIP_2$) systems using Monte Carlo algorithm and MSM transition probability.

**SI Table S1 Dimer configurations in 9-copy systems.** The table is ranked by oligomer residence time and grouped by receptor conformational states. ($\theta_1$, $\theta_2$) is the orientation angles for dimers that describes the relative position of the dimer interface to the principal axis of that monomer that is parallel to H8. A full list of the profiles of oligomer quaternary structures in 9-copy systems can be downloaded at http://doi.org/10.5281/zenodo.4300676 along with the atomistic models of the oligomer structures.

| Conf State | Cluster id | $\theta_1$ mean | $\theta_1$ standard deviation | $\theta_2$ mean | $\theta_2$ standard deviation | Count | Pop. within Oligomeric Order (100%) | Pop. within Conf. State (100%) | Residence Time (µs) |
|---|---|---|---|---|---|---|---|---|---|
| Inactive | 2.9 | 158.1 | 14.055 | 335.68 | 16.69 | 1760 | 6.18 | 4.8 | 14.268 |
| Inactive | 2.0 | 186.57 | 13.608 | 196.55 | 18.33 | 4994 | 17.534 | 13.62 | 10.98 |
| Inactive | 2.8 | 41.26 | 11.348 | 172.92 | 18.8 | 9627 | 33.801 | 26.25 | 8.789 |
| Inactive | 2.2 | 31.539 | 9.326 | 30.041 | 11.01 | 770 | 2.704 | 2.1 | 6.668 |
| Inactive | 2.1 | 45.46 | 88.124 | 179.3 | 151.4 | 3997 | 14.034 | 10.9 | 5.987 |
| Inactive | 2.6 | 42.082 | 58.813 | 242.7 | 31.6 | 1270 | 4.459 | 3.463 | 2.954 |
| Inactive | 2.3 | 96.869 | 32.643 | 164.29 | 22.83 | 2690 | 9.445 | 7.336 | 2.43 |
| Inactive | 2.10 | 28.832 | 30.525 | 127.33 | 61.27 | 388 | 1.362 | 1.058 | 2.131 |
| Inactive | 2.4 | 34.068 | 27.016 | 214.89 | 21.62 | 1383 | 4.856 | 3.771 | 1.446 |
| Inactive | 2.5 | 164.5 | 15.639 | 262.3 | 15.74 | 554 | 1.945 | 1.511 | 1.346 |
| Inactive | 2.7 | 60.735 | 12.125 | 119.58 | 11.48 | 1048 | 3.68 | 2.858 | 0.831 |
| Active | 2.5 | 73.04 | 10.21 | 206.7 | 14.67 | 7588 | 24.723 | 15.59 | 28.523 |
| Active | 2.0 | 120.37 | 19.451 | 153.37 | 22.53 | 7238 | 23.583 | 14.87 | 10.66 |
| Active | 2.6 | 35.107 | 12.243 | 46.137 | 38.7 | 2867 | 9.341 | 5.891 | 9.796 |

| | | | | | | | | | |
|---|---|---|---|---|---|---|---|---|---|
| Active | 2.9 | 96.956 | 9.334 | 114.78 | 10.62 | 3065 | 9.986 | 6.298 | 4.298 |
| Active | 2.7 | 216.1 | 15.216 | 249.09 | 15.18 | 738 | 2.405 | 1.517 | 3.962 |
| Active | 2.4 | 43.788 | 10.18 | 259.36 | 11.67 | 1563 | 5.093 | 3.212 | 3.619 |
| Active | 2.11 | 119.32 | 58.137 | 293.1 | 72.73 | 539 | 1.756 | 1.108 | 3.322 |
| Active | 2.8 | 150.86 | 18.935 | 254.34 | 27.11 | 2276 | 7.416 | 4.677 | 3.232 |
| Active | 2.1 | 67.279 | 20.193 | 138 | 20.02 | 3912 | 12.746 | 8.039 | 2.904 |
| Active | 2.3 | 39.024 | 39.043 | 162.26 | 76.17 | 565 | 1.841 | 1.161 | 2.18 |
| Active | 2.2 | 218.42 | 8.567 | 322.2 | 5.257 | 185 | 0.603 | 0.38 | 1.818 |
| Active | 2.10 | 252.32 | 10.773 | 289.26 | 9.706 | 156 | 0.508 | 0.321 | 1.286 |
| Act + mini Gs | 2.2 | 33.593 | 37.248 | 246.08 | 21.18 | 2706 | 11.642 | 6.001 | 20.038 |
| Act + mini Gs | 2.3 | 88.643 | 10.232 | 99.881 | 6.386 | 2469 | 10.622 | 5.475 | 17.246 |
| Act + mini Gs | 2.8 | 156.66 | 29.031 | 237.3 | 21.21 | 2840 | 12.218 | 6.298 | 13.469 |
| Act + mini Gs | 2.4 | 49.66 | 42.756 | 177.28 | 60.4 | 8032 | 34.555 | 17.81 | 12.181 |
| Act + mini Gs | 2.6 | 27.825 | 6.341 | 41.415 | 25.77 | 1167 | 5.021 | 2.588 | 11.188 |
| Act + mini Gs | 2.0 | 57.736 | 11.759 | 225.47 | 103.2 | 747 | 3.214 | 1.657 | 9.264 |
| Act + mini Gs | 2.1 | 99.162 | 9.353 | 322.22 | 22.61 | 1163 | 5.003 | 2.579 | 7.631 |
| Act + mini Gs | 2.7 | 133.43 | 15.946 | 161.57 | 18.78 | 1793 | 7.714 | 3.976 | 5.22 |
| Act + mini Gs | 2.9 | 192.1 | 22.976 | 268 | 11.22 | 834 | 3.588 | 1.85 | 4.113 |
| Act + mini Gs | 2.5 | 89.807 | 55.375 | 257.91 | 102.5 | 1433 | 6.165 | 3.178 | 3.758 |

**SI Table S2 Dimer configurations in 9-copy NoPIP$_2$ systems.** The table is ranked by oligomer residence time and grouped by receptor conformational states. ($\theta_1$, $\theta_2$) is the orientation angles for dimers that describes the relative position of the dimer interface to the principal axis of that monomer that is parallel to H8.

| Conf State | Cluster ID | $\theta_1$ mean | $\theta_1$ standard deviation | $\theta_2$ mean | $\theta_2$ standard deviation | Count | Pop. within Oligomeric Order (100%) | Pop. within Conf. State (100%) | Residence Time (μs) |
|---|---|---|---|---|---|---|---|---|---|
| Inactive | 2.1 | 122.567 | 143.739 | 304.582 | 104.287 | 2444 | 70.636 | 62.554 | 4.129 |
| Inactive | 2.3 | 30.216 | 36.146 | 198.985 | 28.552 | 665 | 19.22 | 17.021 | 2.857 |
| Inactive | 2.2 | 41.046 | 69.758 | 261.902 | 30.512 | 244 | 7.052 | 6.245 | 2.332 |
| Inactive | 2.4 | 143.189 | 66.26 | 279.784 | 88.525 | 28 | 0.809 | 0.717 | 0.86 |
| Inactive | 2.0 | 195.38 | 22.889 | 239.899 | 19.596 | 73 | 2.11 | 1.868 | 0.515 |
| Inactive | 2.5 | 145.171 | 10.296 | 167.204 | 14.867 | 6 | 0.173 | 0.154 | 0.047 |
| Active | 2.2 | 115.828 | 63.94 | 279.229 | 85.747 | 2022 | 30.306 | 27.634 | 5.82 |
| Active | 2.0 | 36.138 | 9.478 | 56.205 | 17.005 | 1689 | 25.315 | 23.083 | 4.097 |
| Active | 2.5 | 124.762 | 11.296 | 144.383 | 13.628 | 2268 | 33.993 | 30.996 | 5.525 |
| Active | 2.3 | 46.414 | 12.767 | 137.728 | 11.167 | 304 | 4.556 | 4.155 | 2.235 |
| Active | 2.4 | 252.179 | 8.748 | 295.594 | 8.506 | 194 | 2.908 | 2.651 | 1.886 |
| Active | 2.1 | 52.152 | 60.229 | 254.401 | 30.742 | 156 | 2.338 | 2.132 | 1.629 |
| Active | 2.6 | 213.984 | 17.362 | 247.049 | 20.423 | 39 | 0.585 | 0.533 | 0.223 |
| Act + mini Gs | 2.5 | 77.782 | 96.107 | 265.205 | 55.63 | 3052 | 47.755 | 28.42 | 12.108 |

| | | | | | | | | | |
|---|---|---|---|---|---|---|---|---|---|
| Act + mini Gs | 2.2 | 59.922 | 18.823 | 296.279 | 104.887 | 1001 | 15.663 | 9.321 | 8.221 |
| Act + mini Gs | 2.6 | 21.064 | 28.705 | 83.135 | 114.278 | 1109 | 17.353 | 10.327 | 5.331 |
| Act + mini Gs | 2.0 | 97.71 | 7.33 | 170.843 | 19.55 | 384 | 6.008 | 3.576 | 3.891 |
| Act + mini Gs | 2.3 | 13.463 | 4.011 | 348.238 | 35.031 | 186 | 2.91 | 1.732 | 2.931 |
| Act + mini Gs | 2.4 | 43.219 | 107.621 | 262.728 | 153.524 | 327 | 5.117 | 3.045 | 2.194 |
| Act + mini Gs | 2.1 | 221.969 | 21.807 | 264.823 | 13.759 | 289 | 4.522 | 2.691 | 1.133 |
| Act + mini Gs | 2.7 | 28.605 | 26.413 | 119.852 | 104.548 | 24 | 0.376 | 0.223 | 1.081 |
| Act + mini Gs | 2.8 | 209.959 | 7.539 | 225.212 | 6.364 | 19 | 0.297 | 0.177 | 0.3 |

**SI Table S3. Lifetimes of oligomerisation states calculated from MSMs of 9-copy systems calculated.** The MSMs were built with a lagtime of 5 µs, thus lifetimes of shorter than 5 µs cannot be resolved.

| Oligomerisation State | Inactive (µs) | Active (µs) | Act. + mini Gs (µs) |
|---|---|---|---|
| $1^9$ | 8.23 | 6.13 | 5.17 |
| $1^7$-$2^1$ | 8.27 | 7.70 | 5.84 |
| $1^5$-$2^2$ | 7.69 | 6.87 | 7.29 |
| $1^3$-$2^3$ | 8.70 | 7.01 | 7.23 |
| $1^1$-$2^4$ | 5.00 | 6.38 | 5.00 |
| $1^6$-$3^1$ | 5.88 | 6.26 | 7.42 |
| $1^4$-$2^1$-$3^1$ | 7.60 | 6.86 | 7.64 |
| $1^2$-$2^2$-$3^1$ | 7.15 | 8.52 | 7.52 |
| $1^1$-$2^1$-$3^2$ | 5.47 | 8.39 | 5.26 |
| $1^3$-$3^2$ | 5.47 | 7.50 | 5.98 |
| $2^3$-$3^1$ | 5.00 | 5.00 | 5.05 |
| $1^1$-$2^2$-$4^1$ | 5.00 | 5.95 | 8.90 |
| $1^3$-$2^1$-$4^1$ | 5.33 | 6.17 | 8.56 |
| $1^5$-$4^1$ | 5.86 | 5.17 | 5.36 |
| $1^2$-$3^1$-$4^1$ | 5.00 | 7.29 | 14.49 |
| $1^1$-$4^2$ | 5.00 | 5.00 | 5.00 |
| $2^1$-$3^1$-$4^1$ | 5.00 | 5.00 | 5.34 |
| $1^2$-$2^1$-$5^1$ | 5.00 | 8.35 | 7.79 |
| $1^4$-$5^1$ | 5.32 | 6.79 | 14.62 |

| | | | |
|---|---|---|---|
| *1¹-2¹-6¹* | 5.00 | 5.00 | 5.10 |
| *1³-6¹* | 5.00 | 5.09 | 5.21 |
| *2²-5¹* | | 5.00 | 7.45 |
| *1¹-3¹-5¹* | | 8.22 | 6.85 |
| *1²-7¹* | | 5.00 | 5.53 |
| *2¹-7¹* | | 5.00 | 5.00 |
| *1¹-8¹* | | 5.14 | 9.73 |
| *4¹-5¹* | | | 6.25 |
| *3¹-6¹* | | | 5.00 |
| *9¹* | | | 5.00 |

**SI Table S4. Lifetimes of oligomerisation states calculated from MSMs of 9-copy NoPIP$_2$ systems calculated.** The MSMs were built with a lagtime of 5 μs, thus lifetimes of shorter than 5 μs cannot be resolved.

| Oligomerisation State | Inactive | Active | Act. + mini Gs |
|---|---|---|---|
| $1^9$ | 11.47 | 5.16 | |
| $1^7$-$2^1$ | 14.65 | 7.13 | 5.27 |
| $1^5$-$2^2$ | 12.27 | 11.09 | 5.87 |
| $1^4$-$2^1$-$3^1$ | 5.56 | 5.77 | 8.88 |
| $1^6$-$3^1$ | 5.48 | 5.00 | 9.39 |
| $1^5$-$4^1$ | 5.00 | | |
| $1^3$-$2^3$ | | 5.57 | 5.00 |
| $1^2$-$2^2$-$3^1$ | | 5.00 | 5.02 |
| $1^3$-$2^1$-$4^1$ | | 5.00 | 5.00 |
| $1^3$-$3^2$ | | | 5.89 |
| $1^1$-$2^1$-$3^2$ | | | 5.00 |
| $1^2$-$3^1$-$4^1$ | | | 5.00 |
